# Tuning in Sensorimotor Synchronization

**DOI:** 10.1101/2022.12.16.520727

**Authors:** Georgios Michalareas, Matthias Grabenhorst, Yue Sun

**Affiliations:** Max Planck Institute for Empirical Aesthetics, Frankfurt, Germany; Ernst Strüngmann Institute (ESI) for Neuroscience, Frankfurt, Germany

## Abstract

Moving in synchrony to external rhythmic stimuli is an elementary function that humans regularly engage in. It is termed “sensorimotor synchronization” and it is governed by two main parameters, the period and the phase of the movement with respect to the external rhythm. There has been an extensive body of research on the characteristics of these parameters, primarily once the movement synchronization has reached a steady-state level. Particular interest has been shown about how these parameters are corrected when there are deviations for the steady-state level. However, little is known about the initial “tuning-in” interval, when one aligns the movement to the external rhythm from rest. The current work investigates this “tuning-in” period for each of the four limbs and makes various novel contributions in the understanding of sensorimotor synchronization. The results suggest that phase and period alignment appear to be separate processes. Phase alignment involves limb-specific somatosensory memory in the order of minutes while period alignment has very limited memory usage. Phase alignment is the primary task but then the brain switches to period alignment where it spends most its resources. In overall this work suggests a central, cognitive role of period alignment and a peripheral, sensorimotor role of phase alignment.

**Highlights:**

- In the tuning-in phase there are three distinct temporal scales of sensorimotor synchronization with distinct signatures. A long-range, across-blocks monotonic negative gradient to more anticipatory movement, which prevails for tens of minutes, a very consistent “hook”-shaped pattern within each block, in the range of seconds, and a constant difference across time between feet and hands.
- The across-blocks, monotonic, negative gradient to more anticipatory movement is instantiated only in the first anticipatory trial of each block and the rest of the subsequent block trials contribute to the alignment of the inter-movement interval to the metronome’s period.
- This negative asynchrony gradient is limb-specific and is not affected by the interleaved blocks of other limbs.
- Period alignment has a central, cognitive role while phase alignment a peripheral, sensorimotor role.

## Introduction

Tapping in synchrony to a metronome is one of the most predictable and least-demanding tasks for the human brain. More generally, synchronization to an external regular rhythm is typically termed sensorimotor synchronization and musical beat is an intuitive example of such an external regular rhythm to which humans can align their movement with relatively little effort. Although for humans this capacity is considered elementary, in almost all other species it is upsent, with very few but notable exceptions(Bouwer et al., 2021).

Two main factors are used to describe SMS (Repp, 2005; Repp & Su, 2013). The first is the Stimulus-Movement Asynchrony (termed here SMA), which is the time difference between the metronome stimulus onset and the corresponding motor movement that is intended to align to it. In the ideal case when movement is perfectly aligned with the metronome beat, the SMA is zero. The second factor is the Inter-Movement Interval (termed here IMI), which is the time difference between two successive movements. In the ideal case of perfect synchronization to the metronome, IMI is equal to the period of the metronome. Sensorimotor synchronization in reality is usually different from the ideal state described above. IMI represents the period and SMA the phase of the synchronized movement with respect to the metronome.

One of the most interesting and largely unresolved phenomena related to sensorimotor synchronization is the overall negative level of the asynchrony. Negative SMA means that the movement (i.e. finger tap) precedes on average the metronome stimulus, typically by tens of milliseconds. This phenomenon has been typically termed as the Negative Mean Asynchrony (NME)(G. Aschersleben, 2002). As it occurs only with highly regular, and thus highly predictable, sequences of stimuli, it has been attributed to some aspects of the anticipatory mechanisms of the nervous system(Repp, 2005). It has been suggested that due to the higher speed of information conduction through audition (or vision) w.r.t to somatosensation, the brain initiates the movement earlier than the stimulus in order to achieve their coincidence(Fraisse, 1980; Paillard, 1946). Another proposed theory has postulated that it is the process of evidence accumulation from different sensory channels in the brain that is the reason behind the asynchrony between movement and stimulus(G. Aschersleben, 2002; Aschersleben et al., 2004). As tactile information is less accurate than auditory or visual, the evidence accumulation for registering the tap of a finger takes longer and thus the brain starts earlier the movement so that evidence accumulation from the different sensory modalities coincides at a given perceptual threshold. Evidence for this comes from the fact that the asynchrony decreases when the auditory feedback from a tap is enhanced and probably the brain starts using the auditory evidence of the tap rather than the tactile to align it to the metronome(Aschersleben & Prinz, 1995; Aschersleben & Prinz, 1997). Despite these and other proposed theories of the negative bias of SMA, each own with its own merit, none of them has managed to describe adequately all its observed aspects and the causes behind it are still unknown (Repp, 2005).

Regarding the Inter-Movement Interval, IMI, no such systematic bias has been observed and it is assumed that it converges to the actual period of the metronome. However even for IMI there are still some unexplained aspects. One such aspect is the that when one is tapping in synchrony to a metronome and the metronome suddenly changes period with a step change, then the movement aligns to it with an overshoot. When the step change of the period is large then the overshoot is big and the convergence to the new period fast. If the step change is small then there is no overshoot and the convergence to the new rhythm happens slowly with a prolonged slight overestimation of the new period (Michon & van der Valk, 1967; Repp, 2001).There is still no adequate explanation of this phenomenon.

What has been studied extensively and described well for both SMA and IMI is the corrective mechanism that preserves the steady-state overall level of sensorimotor synchronization in a trial-to-trial or more intuitively in a beat-by-beat manner(Repp, 2005; Repp & Su, 2013). The term steady-state refers to the condition when after a small number of trials/beats since the start of the metronome, the synchronizing movement has settled to a stable state, and its two main characteristics, SMA and IMI, have both reached nominal levels. After this convergence, a continuous corrective mechanism realigns these factors back towards their nominal values when there are deviations from them and keeps synchronization in a stable state(Repp, 2005). A multitude of descriptive models have been developed throughout the years trying to capture this corrective mechanism. One family of models assumes that there is one single corrective mechanism for both SMA and IMI which is based on SMA as the primary factor. (Michon & van der Valk, 1967; Pressing, 1998; Pressing & Jolley-Rogers, 1997; Schulze & Vorberg, 2002; Semjen et al., 1998; Vorberg, 1996). Correction of IMI in these models is based on the SMA values of the previous two values. Another family of models treat SMA and IMI as being corrected by different mechanisms, and these are typically called dual-process models (Hary & Moore, 1987; Mates, 1994a, 1994b). The main concept of these models is that the IMI correction is performed by a central timekeeper mechanism in the brain while the phase correction is performed by a separate more peripheral mechanism. Mathematically there is no obvious evidence that one family of models is better than the other and actually they can all be converted to an equivalent general form, which uses SMA from the previous two trials as the primary factor for correcting both SMA and IMI(Jacoby & Repp, 2012). However, there is some behavioral evidence that period and phase correction are two different processes with distinct characteristics. Repp (Repp, 2000) first showed that phase correction in steady-state sensorimotor synchronization does not require awareness of phase changes but it rather happens automatically. He then went on to show that period correction, in contrast to phase, is facilitated by conscious awareness of a tempo change. Finally Repp and Keller (Repp & Keller, 2004) showed that higher cognitive processes such as intention and attention affect period correction but have little effect on phase correction. Based on these results, Repp and colleagues proposed that period correction is more of a cognitive, central brain process while phase correction is more of a sensorimotor, peripheral process. There is clear parallelism of this hypothesis with the model of sensorimotor synchronization by (Wing & Kristofferson, 1973) which distinguishes sources of variance to peripheral motor sources on one hand and central timekeeper sources on the other.

All these descriptive linear models of lag 1 or 2 have demonstrated that after the brain has reached a steady-state sensorimotor synchronization to a metronome it works in an almost beat-by-beat fashion to retain this steady state alignment, without the need for any longer short-term memory of its recent behavior. And although this seems to capture well the behavior of the human brain in the steady-state condition, very little is known about the characteristics of sensorimotor synchronization during the initial tuning-in transition to the steady-state condition from rest.

This tuning-in transition from resting to steady-state is important and interesting for various reasons. The most obvious is to study how fast SMA and IMI converge on average to their steady-state levels and what is the shape and the characteristics of these curves. One of the most important and unknown characteristic is whether in this tuning-in process the priority is given first to the alignment of the asynchrony (SMA) and then to the period (IMI), or vice versa. Or even if the convergence to steady-state involves concurrent alignment of both parameters. Studying the tuning-in phase provides the advantage that the brain starts from rest to adapt the metronome and if there is a priority for one of the two parameters, it could be manifested in the tuning-in curves of these two variables, probably as the one with the priority settling in faster to the steady-state level.

Another interesting question regarding the tuning-in phase of sensorimotor synchronization is whether there are significant differences between different limbs. It is already known that in the steady-state phase of synchronization the feet have consistently higher negative SMA, attributed to either longer conduction times or, more likely, to the different effector movement characteristics (Aschersleben & Prinz, 1995; G. Aschersleben, Stenneken, P., Cole, J., & Prinz, W., 2002; Billon et al., 1996). However, it is not known yet whether feet and hands have different tuning-in curves and whether these curves and the converged steady-state values are affected when intervals of synchronization by one limb are interleaved by segments of synchronization of other limbs.

In the current study we investigated the tuning-in phase of sensorimotor synchronization in all 4 limbs. The study used data from the Human Connectome Project(Van Essen et al., 2013), in which the participants synchronized to a visual metronome (period of 1.2sec) from rest in a long sequence of short blocks, each consisting of 10 trials. At the beginning of each block the participants were instructed to synchronize to the metronome with a specific limb, out of four possible limbs, with the exception of few blocks where there was no metronome and the participant were instructed to rest. The block size was long enough to ensure convergence to steady-state, as typically in the study of steady-state sensorimotor synchronization it is assumed that by the tenth trial the asynchrony has reached its steady state level.

The two main parameters, SMA and IMI, were examined in terms of their mean and standard deviation at two different levels, namely within-block and across-blocks. At the microlevel, within-block, we examined for each limb the average and standard deviation of SMA and IMI in each of the ten trials within the block. Each of these within-block trials will be referred to as “*block-trials*” in the remaining of the manuscript, taking values from 1 to 10 in each block, in order to avoid confusion with the typical notion of the continuously increasing trial number across an experiment. Here the focus was on the characteristics of SMA and IMI during the alignment or “tuning-in” phase to the rhythm of the metronome in each block. At the macrolevel, across-blocks, we examined SMA and IMI for each limb in terms of how the block average and standard deviation, as well as the single block-trials, changed across blocks in the course of the experiment. Here an important aspect of investigation was how consistent or stable are the block statistics of SMA and IMI for a specific limb across blocks. Another important question here was if there is modulation of SMA and IMI by the number of interleaved blocks between two successive blocks of the same limb. These interleaved blocks were mostly blocks of sensorimotor synchronization with other limbs and some resting blocks.

One of the a-priori expectations was that feet would have higher negative asynchrony with respect to hands. This difference has been demonstrated in previous research as mentioned earlier. Another a-priori expectation was that the average asynchrony would remain stable, around the same negative level across the experiment. This expectation was based on previous research that has attributed asynchrony to differences between tactile and auditory/visual modalities in the way that sensory evidence is transmitted to and/or accumulated in the brain(Repp, 2005). These processes are more related to structural and elementary functional characteristics, which are not expected to be altered during the course of the experiment. Finally, another a-priori expectation was that within-block the curves for both asynchrony and IMI would converge to their steady state values faster in the later part of the experiment. This is because it was expected that after extended exposure to the fixed metronome rhythm and the fixed trial structure within each block, the brain would have higher anticipatory power and would reach its steady state target faster.

From the above a-priori expectations only the one about the consistent difference between the SMA of hands and feet was confirmed. In contrast, the average SMA was not only found to be unstable across blocks but strikingly it was found to follow a very prominent monotonic negative gradient for the entirety of the experiment. Also, the within-block curves of asynchrony and IMI did not converge faster to their steady state values in the later part of the experiment but they were similar to the ones from the earlier part. An intriguing novel finding regarding the within-block curves was that only the first anticipatory block-trial contributed to the alignment of SMA to its steady-state level, while all the remaining block-trials contributed mostly to the alignment of IMI to its steady-state level. This is an important finding as it shows that the asynchrony seems to be the parameter that has the first priority in sensorimotor synchronization but that alignment to the metronome’s period takes over quickly in priority and dominates the dedicated resources by the brain. On the top of these surprising results we found strong evidence that SMA is a limb-specific phenomenon without transfer to other limbs. Altogether these novel findings show a complicated interplay between SMA and IMI, more complicated than in the traditional models of steady-state sensorimotor synchronization.

## Methods

### Data

The data used in this study was from 61 young healthy adults (ages 22-35) from the HCP1200 Release (HCP1200, 2017) of the Human Connectome Project (HCP). The data was specifically taken from the Motor Task of the unprocessed Magnetoencephalography (MEG) dataset. This task is described in detail in the HCP1200 Release Reference Manual(HCP1200Manual, 2018). Some of the details from there will be reproduced below in various parts of the Methods section for the convenience of the reader and completeness of the manuscript.

### Experimental Design

The Motor Task of the Human Connectome Project aimed at studying sensory-motor processing using a task of motor synchronization to a visual metronome with a fixed period of 1.2 sec. The experiment was performed in a block-design, where in each block the participants had to synchronize with either the left hand, right hand, left foot or right foot to the metronome. At the beginning of each block a visual cue instructed the participants which limb to synchronize to the metronome. The task was adapted from Buckner and colleagues(Buckner et al.; Yeo et al.). The motor movement that was synchronized to the visual metronome involved flexion of either fingers or toes. Specifically in the case of left or right hand the participants were instructed to tap their index finger and thumb with each other. In the case of feet, the participants were instructed to squeeze their toes. This movements are depicted in Fig.1A.

**Figure 1.**
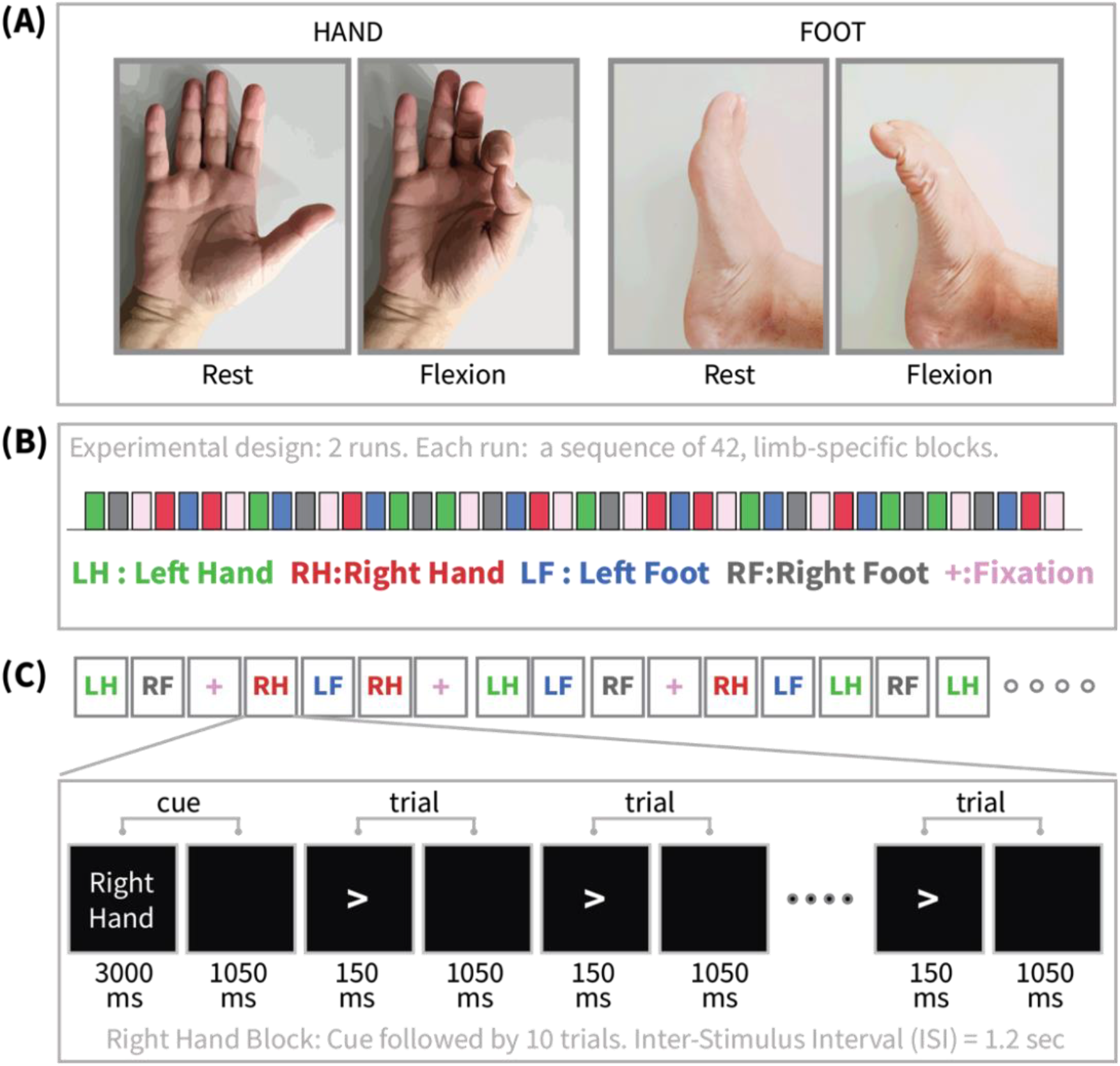
Experiment Description. (A) The participants were asked to move one of their limbs in synchrony with a visual metronome with period of 1.2 seconds. Hand Movement (either left or right): The participants were instructed to tap their index finger and thumb with each other (flexion) from rest. Foot Movement (either left or right): The participants were instructed to squeeze their toes (flexion) from rest. (B) Exact block sequence of a single run. The experiment comprised of 2 identical runs with a break in between. The block sequence was comprised of 42 interleaved blocks, 32 of limb movement and 10 of resting. (C) Structure of a single block. A cue at the beginning instructed the participant which limb to synchronize to the metronome in the current block. Then, there were 10 isochronous stimuli with Inter-Stimulus Interval (ISI) of 1.2 seconds. The stimuli were arrows pointing left or right in accordance with the cue instruction. They were used to assist the participant in not getting confused about which side’s limb they should move.

Each block started with an instruction screen, indicating the side (left, right) and the limb (hand, foot) to be used by the subject in the current block. This instruction screen lasted 3 seconds, followed by a black screen for 1050 msec. Then, 10 pacing stimuli were presented in sequence, each one instructing the participant to make a brisk movement. The pacing stimulus consisted of a small arrow in the center of the screen pointing to the side of the limb movement (left or right). The interval between consecutive stimuli was fixed to 1200 msec. The arrow stayed on the screen for 150 msec and for the remaining 1050 msec the screen was black. Figure 1C shows the structure of a single movement block (In this case in a Right Foot Block).

There were 2 identical runs of the same sequence of blocks. In each run there were 32 limb movement blocks, 16 of hand movement (8 left, 8 right) and 16 of foot movement (8 left, 8 right). In addition to the blocks of limb movements there were 10 interleaved resting blocks, each one of 15 sec duration. During these blocks the screen remained black. The last block was always a resting block after the last limb movement block. As already mentioned, the experiment was performed in identical 2 runs, with a small break between them. In both runs there were 16 blocks (160 trials) per limb and 20 resting blocks. The exact same block sequence of the two runs was presented to all participants. Figure 1B offers a graphic depiction of the block sequence in a single run. The different colors represent different block types.

### Data acquisition

Electromyography (EMG) signals were recorded for capturing muscle activity and identify the onset of hand and foot movement. On the foot EMG sticker electrodes were applied on the lateral superior surface, on the extensor digitorum brevis muscle and near the medial malleolus. On the hand EMG sticker electrodes, also the first dorsal interosseus muscle between thumb and index finger, and the styloid process of the ulna at the wrist. There were two EMG electrodes in each limb and they were recorded in the monopolar form with a common reference, rather than performing the bipolar derivation from them on-the-fly during the recording. This bipolar derivation was performed a posteriori during the analysis.

The EMG channels were part of the MEG recording on a whole head MAGNES 3600 (4D Neuroimaging, San Diego, CA) system housed in a magnetically shielded room, located at the Saint Louis University (SLU) medical campus. The participants were positioned in the MEG scanner in supine position. The electrode impedances of the EMG channels were maintained below 10 kOhms.

Stimuli for the Motor Task were generated using E-Prime 2.0 Pro on a HP personal computer. The stimulus computer presented the visual stimuli through an LCD projector (ImagePro 8935,DUKANE) onto a mirror ∼3 feet above the participant at 1024×768 resolution with 60Hz refresh rate for viewing. Participant button press responses are recorded via fiber optic as the Response channel in the MEG data. Motor-task motion responses, EOG and ECG are recorded via electrodes as described above.

For visual stimuli, the TTL triggers and another digital trigger signal from a photodiode installed on the right-top corner of the screen were combined and recorded as the Trigger channel in the MEG data. All TTL triggers (from the parallel port) of the stimulus computer were recorded as the Trigger channel in the MEG data.

For further details on data acquisition, please refer to the HCP 1200 Release Reference Manual(HCP1200Manual, 2018).

## Ethics

These data were acquired in accordance with the Washington University Institutional Review Board(Van Essen et al., 2013).

### Estimation of SMA and IMI through estimation of Movement Onset from EMG

The SMA and IMI were computed by using the Electromyography (EMG) signals recorded from the corresponding limbs during the experiment. The EMG channels were part of the unprocessed MEG dataset from the Human Connectome Project. Only the EMG channels were extracted from this dataset for each participant. There were 2 monopolar EMG channels for each limb and a bipolar derivation between them was performed at the beginning of the analysis. The resulted bipolar signal will be referred to as “EMG signal” in the rest of the manuscript. The sampling frequency of the EMG signal was *F*_*s*_ = 2034.500 *Hz*.

The electrophysiological correlates of muscle activity, that is related to the movement of interest, are prominent in the upper part of the typical electrophysiology frequency spectrum with the most prominent spectral power in the range 50 to 150 Hz (Bilodeau et al., 1995). So in order to identify the pattern of the motor activity at hand, typically the power envelope of the high-passed EMG filter is used. In detail the following procedure was used.

The EMG movement-onset analysis was performed separately per block. The EMG signal *X* for a given block was extracted and high-pass filtered (using a two-pass 6^th^ order Butterworth IIR filter) with a cutoff frequency of 10Hz. Then the filtered signal was passed through the Hilbert transform which provided its power envelope time-series. Subsequently this time-series was smoothed, in order to remove fast fluctuations, using a moving average filter with length equal to 1/5 of the sampling frequency, which gave a window of 407 samples, corresponding to 0.2 seconds. After this step the resulting processed EMG signal was converted to z-score based on its mean and standard deviation. The analysis up to this point can be termed as “Basic” possessing and typically a threshold close to zero is selected in order to identify the onset of movement. This entails the assumption that the majority of the time there is no movement so the distribution of the processed signal is skewed with the long tail towards the positive values, corresponding to movements. In such case, when the the processed EMG signal is converted to z-score, the main lobe of its distribution is below zero and what is above zero is mostly the long tail corresponding to movement. In such case a threshold of zero makes sense.

In the current experiment though the situation was more complicated and another “Advanced” level of processing was added to tackle some standing issues. The first issue was that the feet movements were much longer than those of the hands and this left less time for baseline (this pushed the mean of the z-score distribution closer to zero). Additionally in various parts of the experiment for each participant there were either short-term or long-term drifts, within some of the blocks, which made the thresholding based on the “Basic” processing problematic. Such short or long-term drifts can be caused by fast or slow movements of the entire arm or leg in parallel to the execution of the hand or foot synchronization respectively. An illustration of these short or long drifts is given in Figures 2A and 2F, where the red lines are the “Basic”-processed EMG signal for a hand and foot block respectively of a single subject. On these figures the movements are depicted as prominent deflections from a baseline of non-movement. A short-term drift that distorts the baseline can be seen at the very beginning of Fig.2A, before the first movement of the hand. There, the “Basic”-processed EMG signal (red) in the first baseline has a much lower level than in the baseline of all the following trials of the block. An example of long-term drift can be seen in “Basic”-processed EMG signal (red) for a block of foot movement in Fig.2F. On this plot it is obvious that there is monotonic negative drift of the baseline. In the cases of both types of drifts, setting up a baseline based on a simple threshold of the “Basic”-processed EMG signal would result in erroneous onset times of movement. For example in the case of the long-term negative drift for the foot in Fig.2F, a single threshold would make the identified movement onset times for the trials in the later part of the block to be in reality actually much later than the actual deflections from the baseline. (Imagine a horizontal line cutting across Fig.2F at a level just below zero).

**Figure 2.**
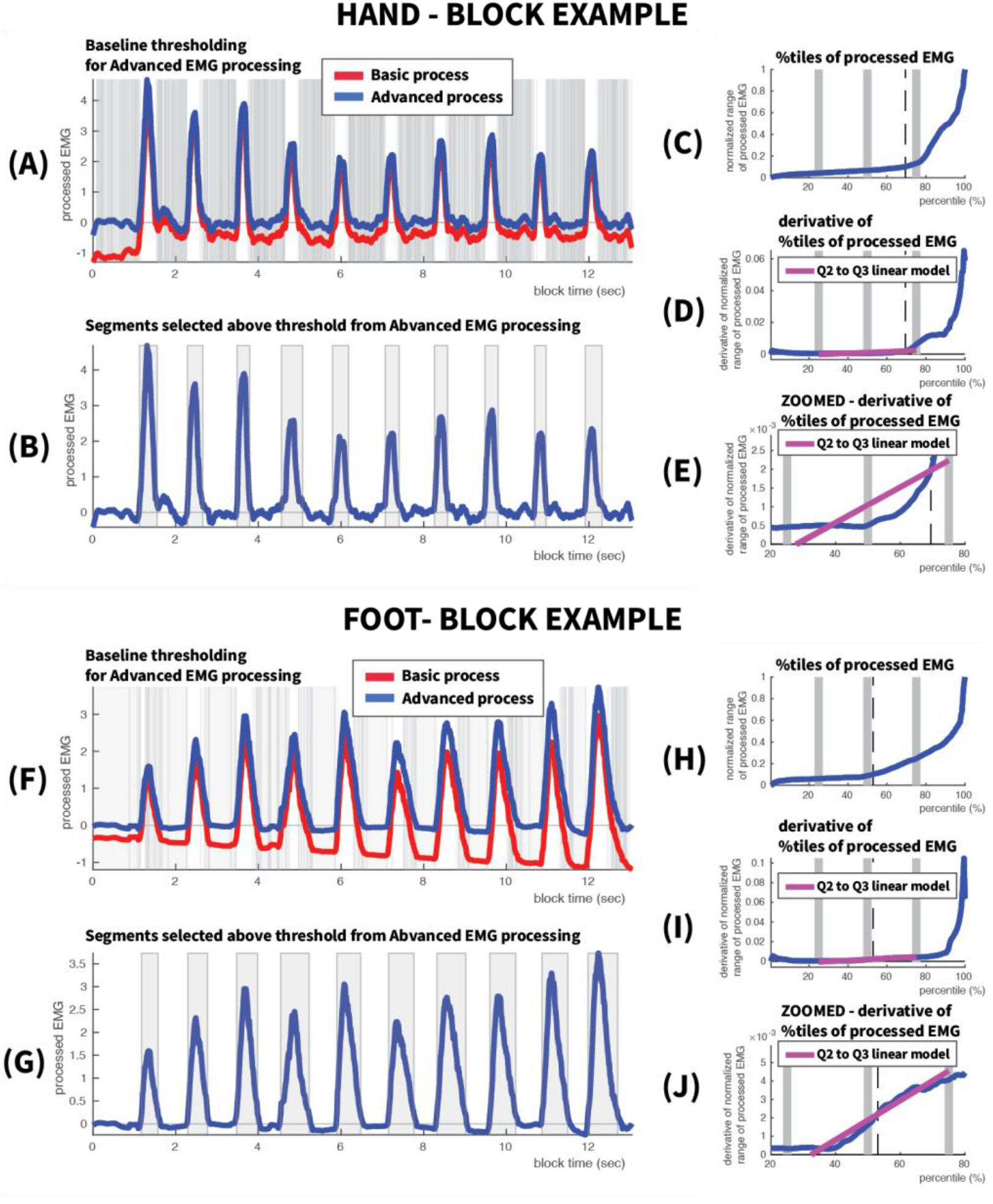
Identification of Movement onset from EMG signal. Single block example for Hand (A)-(E) and Foot (F)-(J). Hand: (A) Processed EMG. Movements correspond to the sharp deflections from baseline. The “Basic” processing (see Methods), depicted in red, suffers from inaccurate baseline identification as well as short-term or long-term drifts. Here such a short drift can be seen in the baseline just before the first movement. The additional “Advanced” processing (see Methods), depicted in blue, removes such drifts and aligns the baseline much better. The thin vertical grey lines correspond to the timepoints used to estimate the baseline in the “Advanced” processing (B) The better baseline alignment and removal of drifts of the “Advanced” processing results in much more accurate selection of movement segments, deflected away from baseline. The beginning of each of these segments was selected as the movement onset (C) Plot of the percentiles of the “Basic”-processed EMG. The thick grey vertical lines are the borders of the quartiles Q1-Q4. (D) The “Advanced” processing uses the derivative of percentiles. It fits a linear function in the derivative mean of quartile Q2 and the mean of quartile Q3. This linear function is depicted with magenta. (E) Same as (D) but zoomed in, so that the linear regression can be seen in more detail. The baseline threshold is selected as the first percentile that crosses over the linear function within quartile Q3. This threshold is depicted by the dashed black line. Feet: (F)-(J). Similar to (A)-(E) but for foot movement. Notice the different pattern of processed EMG deflection corresponding to foot movement in (F) and (G). Foot movement is much more prolonged than hand movement. Notice also the more spread percentile plot for feet in (H) as compared to hands in (C).

In order to avoid these problems and achieve a more appropriate movement onset identification, a further step of “Advanced” analysis was added which removed short-term and long-term drifts. In Figures 2A and 2F the blues lines show the “Advanced”-processed EMG signal, where it can be seen that both short- and long-term drifts have been removed and all trials have comparable baseline before the movement deflections. Based on this cleaner signal the “Advanced” analysis selected a baseline threshold, common for the entire block. The segments that exceeded this threshold and which corresponded to movements are depicted as gray rectangles in Figures 2B and 2G for hands and feet respectively. The successful separation of baseline and movement deflection is apparent for both hands and feet. The start of each of these segments was selected as the movement onset for each trial. Then the SMA was computed by calculating the time difference between this movement onset and the visual metronome stimulus onset of the corresponding trial. Also, the IMI was computed as the time difference between movement onset times of successive trials.

The details of the “Advanced” processing of the EMG signal are the following:

- Let’s call the “Basic”-processed EMG signal, as described above, *Y*
- Also, its derivative across time was computed, which was called *dY*
- The next step was to identify segments of baseline activity, excluding as much as possible movement segments. In order to identify these segments both *Y* and d*Y* were used. Baseline segments of no movement should have relatively low values of EMG signal while at the same time have small changes across time. This was the employed definition of a non-movement segment. In practice this definition was imposed by empirical thresholds on *Y* and d*Y*.
∘The threshold for *Y* was computed as Θ_*Y*_ = *mean*(*Y*) + *std*(*Y*), where *std* stands for standard deviation.
∘Two thresholds were computed for the derivative *dY* (the derivative can be also negative for small movements), a positive (upper) and a negative (lower) threshold as 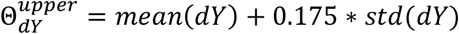 and 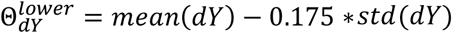 respectively. The weights of the standard deviation here were set empirically after re-iterations across all participants’ data.
- All “Basic”-processed EMG signal data points that satisfied Y < Θ_*Y*_ and 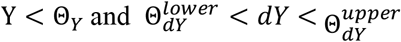 were marked as “baseline-points” with values *Y*_*b*_ and times *t*_*b*_. These points are depicted with vertical, light gray lines in figures Fig. 2A and Fig. 2F for hand and foot respectively. Their correspondence to periods of rest or relatively small movement is apparent.
- Then a polynomial function of 9^th^ degree was fitted on the baseline dataset [*Y*_*b*_, *t*_*b*_]. This function was used to capture any short-term or long-term drifts in the baseline, which should be removed from the data (detrending) before identification of the movement onset. The high degree of the polynomial was selected because there are 10 trials and, as there can be short-term drifts in only one or two trials or long-term drifts across the entire block, the detrending function should be flexible enough to capture both of these types of drifts.
- The polynomial function was evaluated for all the data timepoints, not just those of the “baseline” ones, producing an estimated baseline time-series for the entire block. This estimated baseline time-series was then removed from the “Basic”-processed EMG signal *Y*, leading to the detrended signal *Y*_*D*_.
- The next step was to select the threshold of *Y*_*D*_, above which movement would be defined. Due to the different distribution of *Y*_*D*_ for hands and feet, an empirical heuristic rule was used to compute this threshold for each block separately. This was performed in the following way.
- The percentiles values *P*_*D*_ of *Y*_*D*_ in the range 0.5 to 100 with a step of 0.5 (200 percentile values) were computed and then they were normalized so that they ranged between 0 to 1. These percentiles are presented in Fig, 2C and Fig.2H for hands and feet respectively (blue curves). As it can be seen, the baseline seems to be nicely aligned around zero up to somewhere around 75% for the hand and above 50% for the foot, after which the signal appears to increase rapidly, an obvious manifestation of movement deflections (thick gray vertical lines represent the borders between quartiles Q1:0-25%, Q2:25-50%, Q3:50-75%, Q4:75-100%).
- To quantify this rapid increase in higher part of the percentiles also the derivative of the percentile values across percentiles was computed as *dP*_*D*_. This derivative is depicted in Fig. 2D and Fig.2I for and hand and foot respectively.
- The percentile plot was divided in 4 quartiles (Q1: 0-0.25, Q2: 0.25-0.5, Q3: 0.5-0.75, Q4: 0.75-1). The first quartile Q1 represents the lowest levels of activations which were designated as baseline activity. The upper quartile Q4 was designated as movement activity. The main aim was to identify somewhere within quartile Q3 a threshold which would take also under consideration the rate of increase between quartile Q2 and Q3, as it was identified that the rapid increase in percentile values occur always somewhere in this interval.
- In each of the quartiles Q2 and Q3 the mean derivative *dP*_*D*_ was computed leading to the mean values *μ*_*Q*2_ and *μ*_*Q*3_. These values were assigned to the center of the corresponding quartiles,, i.e. 0.375 and 0.625 respectively, and a first-degree polynomial was fitted on them capturing the slope between the averages of quartile Q3 and Q4. These linear models were extrapolated up to the end of quartile Q3. They are shown in Fig. 2D and Fig.2I with magenta color. As due to the scaling of the percentiles, the slope of the lines and the data they capture appears compressed, a zoomed version of the percentile curves and the linear models is shown in Fig. 2E and Fig 2J for hand and foot respectively. There the different patterns for hands and feet can be clearly seen.
- Then the threshold was set as the point where the normalized percentile values in quartile Q3 crossed above the linear slope. These points are depicted by vertical dashed lines in Fig. 2D,E and Fig.2I,J for hand and foot respectively. For hands the threshold was found more towards the end of Q3 while for feet it was more towards the beginning of Q3. This was expected as the feet movement were longer than for hands (Fig. 2A and 2F) and thus the distribution of the processed EMG signal was more spread across the percentile range.

The above “Advanced” analysis was applied to each block of each participant and from the identified movement onset times the SMA and IMI were computed.

## Results

### Stimulus-Motor Asynchrony (SMA)

SMA was computed for each trial as the time difference between movement onset and the onset of the visual metronome stimulus. At first, the SMA series was examined for the entire duration of the experiment. For each participant all the blocks for each limb were concatenated. This resulted in a series of 160 trials for each limb (16 blocks with 10 trials each). Each of these series was averaged across all subjects, providing the average SMA series. These series for all limbs are presented together in Figure 3A.

**Figure 3.**
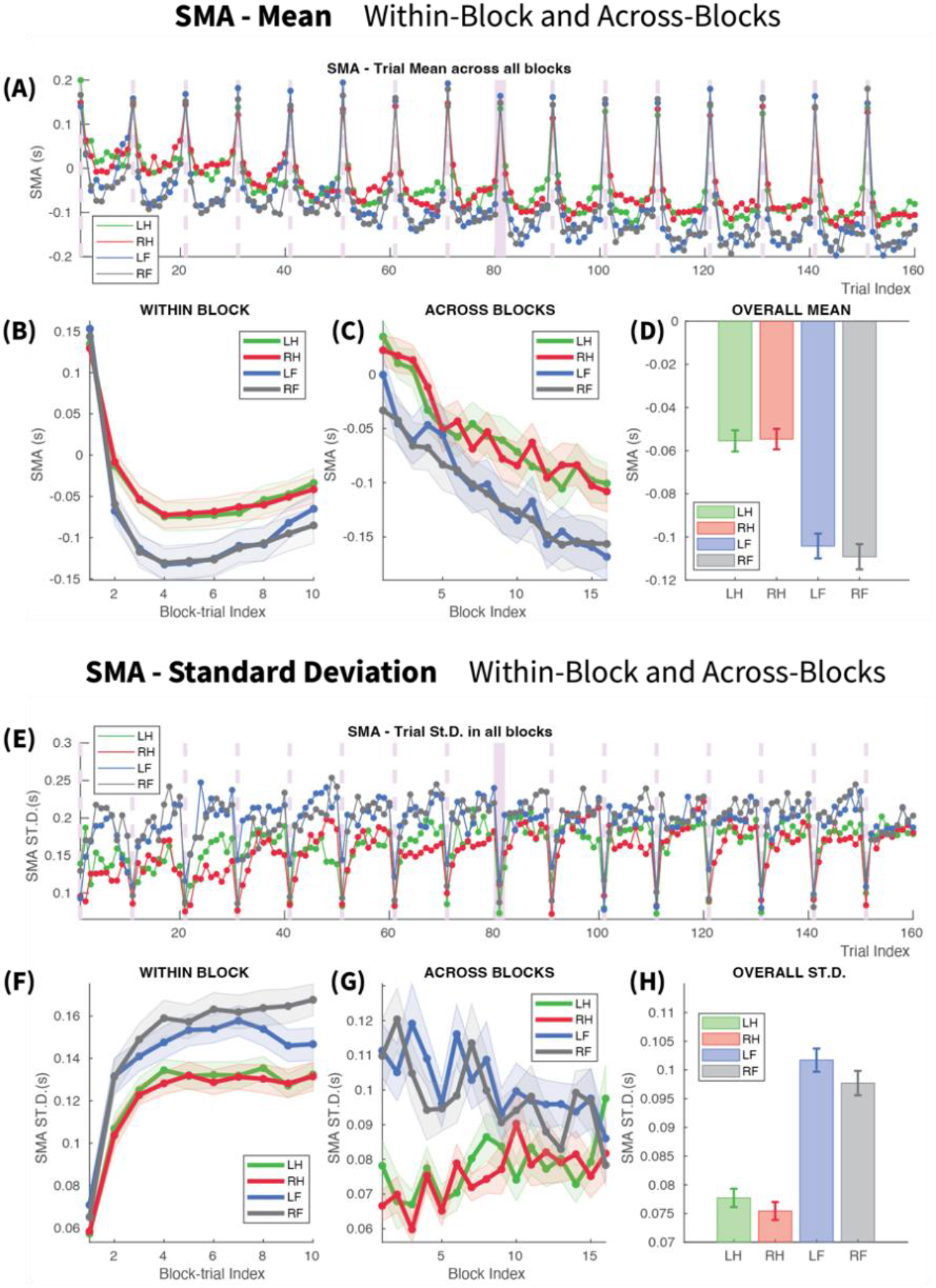
Stimulus-Movement Asynchrony (SMA). (A)-(D): **Study of the SMA Mean.** (A) SMA for each limb in all trials across the entire experiment. (LH,RH: Left/Right Hand, LF,RF: Left/Right Foot). The light purple vertical dashed lines mark the onset of each block. There is a repeated “hook”-shape in each block. The SMA in the first trial of each block is always positive around 150 msec. It is the reaction time to the first metronome stimulus of each block. The rest of the trials show a systematic negative slope across the entire experiment. (B) Within-block SMA. The “hook”-shape. In the first block-trial SMA is positive as it is reaction time. Then from trial 2 onward becomes negative, i.e. anticipatory. The SMA decreases up to trial 4 and then start increasing again. The feet have consistently more negative SMA. (C) Across-blocks SMA. For all limbs there is a monotonic negative gradient of the block average of SMA across the entire experiment. Compare this macrolevel monotonic decrease to the non-monotonic decrease of the within-block curves. (D) Overall mean SMA for the different limbs. The feet have significantly more negative SMA than hands. The difference is −50 msec. (E)-(H): **Study of the SMA Standard Deviation(St.D)**. (E) The St.D of SMA for each limb in all trials across the entire experiment. There is a repeated increasing concave pattern in each block. (F) Within-Block behavior of SMA St.D. The increasing concave pattern of the St.D is seen here clearly. The first block-trial has the smallest St.D. due to the fact that SMA there is reaction time, always around the same value. Then in block-trial 2 SMA has intermediate ST.D and by block-trials 3 and 4 it reaches a maximum plateau and remain there until the end of the block. (G) Across-Blocks behavior of SMA St.D. At the beginning of the experiment Feet have much higher St.D than hands. They then converge across the entire experiment to an intermediate level. Feet get more consistent and hands become less consistent. (H) Overall behavior of SMA St.D. Feet have significantly higher St.D than hands. No significant differences were found between left and right hands of Feet

In these series there are various obvious patterns of dynamical behaviour. The first obvious pattern is the consistent U-shaped periodicity in SMA variation. This pattern is similar in all blocks and for all limbs. The first trial of each block has always a high positive SMA as the participants waits for this trial to start tapping and consequently the SMA in this trial is just reaction time. These firsts trials are clearly visible on the figure and they mark the onset of each block. The second pattern that is obvious is a consistent monotonic negative gradient across blocks. The SMA in the first block starts in most of the trials in the vicinity of 0 sec and by block 16 it has reached a level in the vicinity of −0.2 sec. The third pattern that is obvious in the SMA series is the consistent difference between hands and feet. Feet appear in every block to lag behind the hands, as SMA is consistently more negative for feet. The consistency of these 3 different patterns in SMA across the entire experiment dictated their further investigation.

### Within-block SMA

Within each block the SMA appears to follow a kind of “hook”-like shape. To study this curve, each of the 10 block-trials was averaged across all blocks of a specific limb, providing its average within-block curve per subject. Then, the mean and standard error of the mean (SE) across all subjects were estimated for each block-trial. Figure 3B presents the mean curve across subjects for each limb. The standard error of the mean is represented by the corresponding shaded areas around the mean curves.

The first block-trial, with SMA in the vicinity of 0.15sec for all limbs, is the reaction time to the first metronome stimulus of the block. As such it has a very small standard error across subjects and interestingly it is very similar between all limbs. Another interesting observation about the first trial of the block, from Figure 3A, is that across blocks the SMA remains relatively constant and does not get larger or smaller.

The second block-trial, as it evident from Figure 3B, has already a negative SMA. This means that the participants after the reaction to the first metronome stimulus of the block they immediately deploy their anticipatory model for predicting the second metronome stimulus and this anticipation is manifested as the negative SMA.

This SMA continues to grow more negative in trial 3 and reaches its most negative value in trial 4. After this point, from trial 5 onwards to trial 10 at the end of the block, it retracts monotonically to less negative levels. This monotonic positive retraction after trial 4 (the maximum negative SMA) until the end of the block is a surprising phenomenon. Based on previous research on finger tapping one would expect that once a target SMA has been reached within the first 4-5 trials then in the rest of the trials the participant would just make positive and negative corrections around this level in order to maintain it. Here, it is evident from Figure 2B that this is not the case, but instead SMA has a monotonic positive drift towards 0. This behaviour was identical for all 4 limbs.

The main difference between limbs is that the feet showed consistently larger negative SMA than the hands throughout the block. The only trial that this did not hold was the first trial of each block where no anticipation was present. Beyond trial 1 in each block, the difference between hands and feet appears to be consistent in all other block-trials. In block-trial 3 it is around 60 msec and gradually it is reduced to 40 msec by trial 10. This gradual decrease in SMA difference across block-trials seem to be due to the qualitative difference of curve shapes between hands and feet, apparent in Figure 2B, and could be attributed to different anticipatory mechanisms for different limb systems(hands/feet). This is reinforced by the fact that SMA curves are identical for the two hands and feet respectively.

In order to check whether there was a learning effect on the behavior of SMA we splitted the block sequence in two halves, early (run 1) and late (run 2), and we examined whether in the second part of the experiment the participants’ SMA reached its peak earlier than trial 4. No difference was found between the two halves as seen in Figures 5A and 5B. So the within-block SMA does not appear to benefit from longer exposure to the constant properties of the stimuli.

### Across-Blocks SMA

In order to get a first glimpse of the anticipatory behavior of SMA across blocks, the mean SMA of each block was computed after the first trial of each block was removed, as it corresponded to reaction time. This provided the average anticipatory SMA per block. These across-block curves were derived for each subject and limb. Figure 3C shows these across-block curves, averaged across participants for each limb. It is obvious that the SMA has a similar negative slope across blocks for all limbs. The feet curves have consistently more negative SMA than hands. No lateralization was observed between left and right limbs.

The almost linear slope of SMA across blocks is a surprising finding. There is no obvious reason why the SMA would continue to become steadily more negative across blocks for such a long time interval. If all the blocks for a specific limb had been presented as a contiguous sequence this interval would be equal to the length of 16 blocks, in the order of 4 minutes in total. One would expect that in such a simple metronome task, the SMA would converge to a preferred steady-state level within a few trials (Fraisse & Repp, 2012; Semjen et al., 1998) in the first block of a specific limb and then in each subsequent block of the same limb the SMA would converge within the first few trials in overall to this same steady-state SMA.level. Here in the data of the current experiment no such steady state SMA is evident. SMA keeps becoming more negative across blocks for the entire duration of the experiment.

Even more interestingly, in the current experiment the blocks of the same limb were not presented in a contiguous sequence, but they were rather interleaved with blocks of other limbs and resting blocks. This simply means that between two successive blocks of the same limb there was a number of blocks of other limbs or resting and that the 16 blocks of a single limb were actually presented across a time interval in the order of 20 minutes (see Methods). This is depicted in Figure 1B, where the coloured rectangle sequence demonstrates the actual sequence of blocks, with color representing the kind of limb or resting state. It is obvious that there is a variable number as well as different patterns of interleaved blocks between successive blocks of the same limb. This means that the SMA, which shows a near-constant linear negative gradient across blocks, appears to do so irrespective of the number and pattern of interleaved blocks. Intuitively, between two successive data points in Figure 3C the x-axis step interval is not constant, but varies according to the varying number of interleaved blocks. The persistence of the negative gradient even when blocks of other limbs are interleaved means that there is some form of memory which underlies the gradual descent of SMA across blocks. This across-blocks complicated behavior of a stepwise negative gradient with some form of long short-term memory across blocks is a novel observation. Naturally, follows the question whether this newly observed memory is limb-specific or there is some form of transfer across and interaction between the different limbs. This intriguing question is treated in detail in a later section of this manuscript.

Comparing the across-block with the within-block SMA curves, in Figures 3B and 3C respectively, reveals a perplexing contradiction. The across-block SMA pattern in the macrolevel shows a clear monotonic negatve slope across blocks, while the within-block pattern in the microlevel decreases monotonically only up to trial 4 and then retracts to a positive slope, making a “hook”-like shape. Had the main focus of the brain been to align its SMA to a latent target asynchrony, i.e the SMA of the last block, then it should be expected that also within each block the SMA should monotonically decrease after the first trial, towards this target. The fact that it doesn’t and that it turns upwards after block-trial 4 is evidence that these two patterns might be describing two different mechanisms or functions in somatosensory aniticipation. One could argue that once a block starts, the participant uses only a relatively small number of trials to reach a level of SMA and from this point on the brain switches its focus to another optimization target, which could be a simple homestatic stabilization of SMA based on the immediate few preceding trials or possibly aligning and maintaining a stable inter-movement interval as close as possible to the metronome’s period. (This is treated in detail in the upcoming sections of the manuscript).

### Consistent SMA difference between Hands and Feet

An important aspect that needs to be highlighted here is the range of the negative mean asynchrony for hands and feet and their difference. In Figure 3C it is demonstrated that hands start in the first block with a positive SMA of about 28 msec and gradually across blocks becomes more negative, reaching in the last block an asynchrony of about −104 msec. The initial SMA of 28 msec in the first bock is far smaller from the range of normal reaction times, so it cannot be argued based on this value that in the first block of each limb the participants are just reacting to the stimulus and are not anticipating. Such small positive asynchronies are still considered anticipatory as the average reaction times are significantly higher around 190 msec for visual stimuli and 160 msec for auditory stimuli(Welford & Brebner, 1980). The SMA in the last block, −104 msec, is in the lower end of the typical SMA range of finger tapping experiments, which typically is −20 to −100 msec (G. Aschersleben, 2002). The hand movement in the current experiment is more complex than simple finger tapping. The participants were instructed to flex their thumb and index fingers so that they tap on each other in opposition (see Figure 1A). This movement includes coordination of two different groups of muscles, whose synchronization defines the tapping event itself. These additional levels of motor and sensory complexity and uncertainty make evidence accumulation slower and more variable than for a simple finger tap. In this sense an SMA of −104 msec is still within a range justified by the complexity of the underlying movement. For feet the SMA was consistently more negative than hands as demonstrated in Figure 3B. This was expected as it is already known that toe tapping has more negative SMA than finger tapping, with a difference ranging between −40 to −60 msec (Aschersleben & Prinz, 1995). Consistent with this expected range of differences from hands, in the first block the average feet SMA was around −17msec and became more negative gradually until the last block, where it reached a value of about −162 msec. Here again the foot movement is much more complex that simple toe tapping, involving dorsiflexion and plantarflexion, without a clear tapping target (surface or button). So the kinematics, kinesthetics and the evidence accumulation in the brain for such a complex movement is expected to be slower than simple toe tapping.

In order to quantify the overall difference between hands and feet, the overall mean SMA was computed for each limb. This was performed by first computing for each participant the mean SMA across the entire experiment for each limb. Then for each limb the overall mean SMA was computed across subjects together with the corresponding Standard Error of the Mean. These statistics are presented for each limb in Figure 3D. The mean SMA values for individual subjects were used for performing Wilcoxon signed-rank tests between all combinations of limb pairs. The Wilcoxon test was chosen because the distribution of the mean SMA for all subjects was found to not resemble a normal distribution for any limb, as indicated by the Chi-Square test for normality.

According to the Wilcoxon test, no significant differences were found between Left and Right Hands (LH-RH: Z=-0.427, p<0.6691061) and between Left and Right Feet (LF-RF: Z=1.516, p<0.1293954). In contrast all comparisons between Hands and Feet were found significant (LF-RF: Z=1.516, p<0.1293954, LH-LF: Z=4.218, p<0.0000246, RH-RF: Z=4.572, p<0.0000048, LH-RF: Z=4.478, p<0.0000075, RH-LF: Z=4.446, p<0.0000087).

The overall mean for hands was −55 msec and for feet −105 msec. This means that the flexion onset for the feet occurred about 50 msec earlier than the flexion onset for the hands, with respect to the metronome. This larger negative SMA for feet, with a difference from hands in the range −40 to −60 msec, is a well known phenomenon(Aschersleben & Prinz, 1995). This higher negative asynchrony has been attributed not due to an increased anticipation per se in the brain for the feet, but mostly to the fact that the feet have very different kinematics from hands.

### Variability of SMA

Another important aspect of SMA is its variability. Here in order to study it we computed the standard deviation of SMA for each limb. Figure 3E presents the average standard deviation curves across the entire length of the experiment for all limbs. There is an obvious pattern repetition in each block resembling a concave function, starting at a low value, increasing sharply and then settling at a plateau.

#### Within-block behavior of Standard Deviation of SMA

The within-block curve of the standard deviation was computed by first deriving for each subject and limb the standard deviation across blocks for each of the 10 block-trials. Then these single subject curves were averaged across subjects providing the average standard deviation across subjects and its corresponding standard error(Harding et al., 2014). This average within-block standard deviation curve is shown in Figure 3F for each limb. All limbs have a concave curve. The first trial in the block has the smallest standard deviation, which is expected as there is no anticipation in the first trial, only reaction to the first metronome stimulus. This simple reaction time is similar across all blocks, and this is translated in the first trial of the block having the smallest standard deviation and standard error.

Interestingly also the SMA in the second trial of the block has distinctly small standard deviation, although it is the first trial where movement is anticipatory. After that, trial 3 leads to a plauteau of standard deviation around which revolve also all subsequent trials in a block. There seems to be no obvious, trivial reason why trial 2 in a block has such a smaller standard deviation as compared to trials 3-10 in a block, as the movement there is also anticipatory. Actually trial 2 is the first anticipatory trial in a block.

#### Across-block behavior of SMA Standard deviation

The across-block curve of standard deviation was computed by first computing the standard deviation of each block. Trial 1 from each block was excluded from the computation of its standard deviation due to the fact that SMA there is a reaction time, more or less stable across blocks, while the rest of the trials are anticipatory and have a negative near-linear evolution across blocks. The single-subject curves of standard deviation across blocks were subsequently averaged across subjects, providing the average across-block standard deviation curve and corresponding standard error for each limb. These curves are shown in Figure 3G. There are two clear opposite patterns for hands and feet. For the hands the standard deviation of SMA starts with low values in the first blocks and progressively increases to higher values. For the feet SMA has high standard deviation at the beginning of the experiment and then it progressively has less and less variability. All limbs seem to converge to a similar standard deviation level at the end of the experiment. The patterns for left and right hands are identical and the same holds for feet. This is an interesting pattern very likely reflecting the higher variability in the movement of feet, due to being effectors requiring not so fine movement control as hands. So at the beginning of the experiment this would explain the feet having much higher standard deviation than the hands. As the experiment progressed the feet had less variability in their movement within block, which possibly means that there were less large amplitude noisy fluctuations in the single-subject SMA curves towards the end of the experiment. The participants towards the end of the experiment had performed the same movement many times and this can have caused the refinement of the initially coarse motor control of the movement. However, the unexpected phenomenon was the steady gradual increase of variability in the SMA of hands. It is possible that the fingers, as effectors with refined motor control, start the experiment with the smallest variability but as the experiment progresses and due to its repetitive character the brain makes the motor control progressively coarser as, even so, it is adequate for the task at hand. As the variability of hands and feet converge to an intermediate level towards the end of the experiment, it could be assumed that this is the level of variability and motor control that is adequate for performing this task with any of the four limbs.

#### SMA Standard Deviation difference between Hands and Feet

In both the within-block and across-block behavior of the Standard Deviation of SMA (Fig3E,F,G) it is clearly seen that there is consistently a difference between feet and hands, which towards the end of the experiment becomes progressively smaller, as seen in Figure 3G. The standard deviation is in overall higher for feet. The average standard deviation of SMA was computed for each limb in order to examine the differences between limbs. This was performed by first computing for each participant the standard deviation of SMA across the entire experiment for each limb. Then the average standard deviation was computed across subjects together with its Standard Error. These are shown for each limb in Figure 3H. The values for individual subjects were used for performing Wilcoxon signed-rank tests between all combinations of limb pairs similar to the tests for the SMA.

According to the Wilcoxon test, no significant differences were found between Left and Right Hands (LH-RH: Z=1.246, p<0.2126867) and between Left and Right Feet (LF-RF: 2.204, p<0.0275237). In contrast all comparisons between Hands and Feet were found significant (LF-RF: Z=-4.373, p<0.0000123, LH-LF: Z=-5.808, p<0.0000000, RH-RF: Z=-4.520, p<0.0000062, LH-RF: Z=4.478, p<0.0000075, RH-LF: Z=-5.875, p<0.0000001).

### Summary of results for SMA and its variability

Three main distinct temporal patterns were observed in the SMA of hands and feet:

1. Across blocks, in the long range of multiple minutes, the asynchrony steadily became more negative across blocks with a near-linear slope. This long range slow evolution across blocks is evident of some form of sensorimotor memory, used as a buffer of SMA evolution across blocks.
2. Within block, in the short range of 12 seconds (10 trial of 1.2 seconds each), surprisingly the asynchrony is not becoming more negative monotonically. This appears to occur only for the first 4 trials. After that point for the remaining 6 trials the asynchrony turns upward with a slow positive slope.
3. Within and across blocks there is in overall a consistent negative difference of the order of about −50 msec between hands and feet. This near-fixed difference is agreement with previous literature that has attributed it to different effector kinematics and different sensory accumulator models in the brain for hands and feet.

The fact that within a block the SMA does not continue to become more negative after 4 trials but is reversed, while it has a steady monotonic negative slope across all blocks is strong evidence that there is more than one process that affect the evolution of SMA. Before further speculation on the nature of these multiple processes it is necessary to study the behavior of inter-movement interval (IMI) within- and across-blocks with respect to the metronome period.

The analysis of the standard deviation of SMA revealed three main phenomena:

1. Feet have on overall higher variability in their SMA than hands.
2. The SMA of feet in the first blocks has high variability which steadily decreases until the last blocks. The opposite happens for hands which start with low variability which progressively increases. The variability of hands and feet converge to an intermediate, comparable level towards the end of the experiment and this could be assumed that is the level of variability and motor control that is adequate for performing this task with any of the four limbs.
3. Block-trial 2, the first anticipatory trial, has less variability than the other anticipatory trials, block-trials 3 to 10.

### INTER-MOVEMENT INTERVAL (IMI)

The second and equally important parameter in sensorimotor synchronization is the time interval between two successive movements, termed here Inter-Movement Interval (IMI), which in the absence of any source of stochasticity should be equal to the metronome period, Tm =1.2 sec.

In order to investigate IMI in the current dataset we computed in each trial the time difference between the corresponding movement and the one in the previous trial. Figure 4A shows these IMI values for each trial of each limb across the entire experiment. The first obvious observation is that there is a repeated concave pattern in each block, starting from small IMI values and sharpy converging to a higher level where it settles. This pattern is very similar for all limbs.

**Figure 4.**
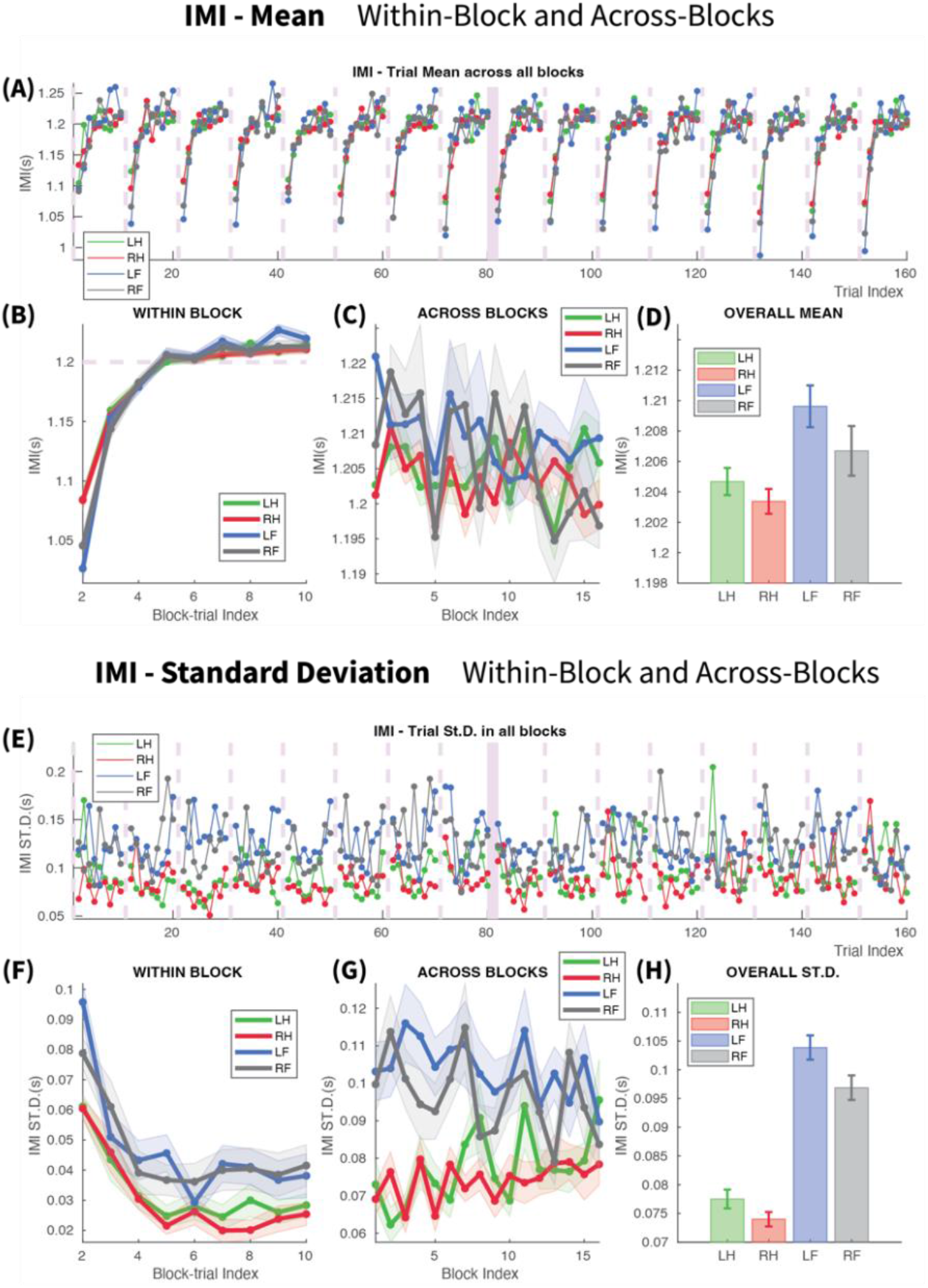
Inter-Movement Interval (IMI). (A)-(D): **Study of the IMI Mean.** (A) IMI for each limb in all trials across the entire experiment. (LH,RH: Left/Right Hand, LF,RF: Left/Right Foot). The light purple vertical dashed lines mark the onset of each block. There is a repeated concave-shape in each block. (B) Within-block IMI. The increasing concave curve is seen clearly here. The first IMI values, corresponding to block-trial 2, is much smaller than the period of the metronome, 1.2 sec, which is shown by a horizontal dashed line. This happens because the SMA in block-trial 1 is positive (reaction time) and in block-trial 2 negative (anticipatory). So the first IMI between them is much shorter than the period. IMI then increases gradually and reaches the vicinity of the metronome’s period in block-trial 5. In block-trials 6 to 10 the IMI settles in a plateau slightly higher than the period. (C) Across-blocks IMI. For all limbs the IMI stays overall slightly above the metronome’s period. (D) Overall mean IMI for the different limbs. They are all above the metronome’s period by a few milliseconds. The only significant differences that were found were Left Foot-Right Foot and Left Foot – Right Hand. (E)-(H): **Study of the IMI Standard Deviation**(St.D). (E) The St.D of IMI for each limb in all trials across the entire experiment. (F) Within-Block behavior of IMI St.D. It follows a decreasing concave pattern. The first block-trial has the highest St.D. and then the St.D gradually decreases until trials4-5 where it reaches a minimum plateau and remains there until the end of the block. (G) Across-Blocks behavior of IMI St.D. At the beginning of the experiment the IMI of Feet has much higher St.D than hands. They then converge across the entire experiment to an intermediate level. Feet get more consistent and hands become less consistent. (H) Overall behavior of IMI St.D. Feet have significantly higher IMI St.D than hands. The IMI St.D was also significantly different between Left and Right Foot. No significant differences were found between left and right hands.

#### Within-block IMI

In order to study the within-block behavior of IMI, first the single-subject within-block curves were calculated by averaging each block-trial across blocks. Then these single-subject curves were averaged across subjects providing the overall mean within-block IMI curve and the corresponding standard error. Figure 4B shows these curves for all limbs. No data is shown for trial 1 in this figure, as IMI is not defined for trial 1. In these plots it is clearly seen that IMI follows in each block an increasing concave curve, starting in trial 2 significantly smaller than the metronome’s period, then reaching with a steep gradient the level of the metronome period at trial 5 and thereafter slightly increasing with a very slow slope until the end of the block. The small IMI value in block-trial 2 is due to the fact that the movement in the first block-trial is always a reaction time, occurring much later than the metronome, while in block-trial 2 the movement is already anticipatory and happens either before the second metronome or slightly after. So it was expected that the time difference between the movements in trial 2 and trial 1 would be much smaller than the metronome’s period. Beyond this trivial behavior of the first IMI in each block, there is a number of other interesting phenomena whose explanation does not appear to be so trivial.

The first of those phenomena has to do with the number of trials that it takes for the concave IMI curve to reach the level of the metronome’s period. It is clearly seen in Figure 4B that the second IMI derived from trials 3 and 2 (plotted on trial 3 of x-axis) is also much smaller than the metronome’s period and the same holds also for the third IMI derived from trials 4 and 3 (plotted on trial 4 of x-axis). So the question arises why does the brain need in each block 5 out 10 trials in order to reach the metronome’s period, when this period has no stochasticity at all and it is just been repeated so many times. The brain should be able to learn and model accurately the metronome s interval within the first few blocks and should be able thereafter to reproduce it faster than the 5^th^ trial.

By splitting the block sequence in two halves, early (run 1) and late (run 2), we examined whether in the later part of the experiment the participants’ IMI reached earlier than trial 5 the metronome ‘s period. This would be evidence that the slow transition in the early part of the experiment would be due to an inadequate capturing of the metronome’s period by the brain which, after longer exposure, became better in the second part of the experiment.. However, no difference was found between the two halves as seen in Figures 5C and 5D. This means that this gradual slow transition to the metronome’s period is a static process, not affected by the amount of exposure time to the metronome, i.e. it is not affected by learning the statistics of the stimulus. A possible explanation for the long transition of IMI to the metronome’s period is that the brain at the beginning of each block is mostly concerned with SMA alignment. Once an SMA level has been reached, the brain switches its focus on aligning its IMI as close as possible to the metronome’s period.

**Figure 5.**
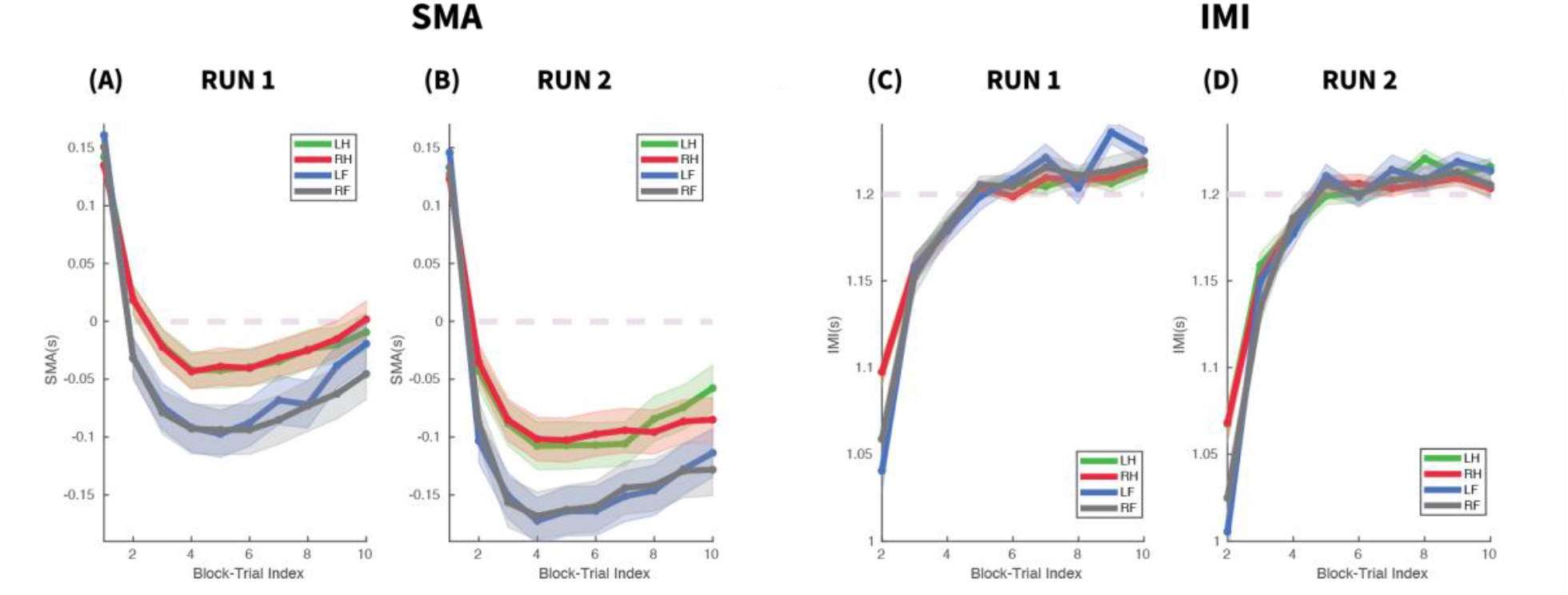
Comparison of Within-Block curves of SMA and IMI between first and second halves of the experiment. (LH,RH: Left/Right Hand, LF,RF: Left/Right Foot). (A),(B): SMA converges to its minimal value in block-trial 4 in both halves. It does not converge faster in the second half. The overall SMA in the second half is much larger (more negative) than in the first half. This was expected due to the negative gradient of SMA across blocks. (C),(D): IMI converges to metronome’s period and reaches it by block-trial 5. It does not converge faster in the second half. After that in block-trials 6 to 10 t remains slightly over the metronome’s period. This overestimation is significant in both halves. No significant difference of this overestimation was found between the two halves.

The second interesting phenomenon observed in the within-block IMI in Figure 3B, is that in trials 7 to 10, where the concave curve seems to be reaching a plateau, the actual IMI level is larger than the metronome’s period Tm =1.2 sec. This appears to be the case for all limbs.

A Wilcoxon test was performed per limb on the IMI difference from 1.2 sec, the metronome’s period. The IMI used in this test was the average of IMI for trials 7 to 10 where the overestimation of metronome period is observed. The test showed that this overestimation was highly significant for all limbs. (LH: Median=1.21072, Z=4.962, p<0.0000007, RH: Median=1.20979, Z=4.468, p<0.0000079, LF: Median=1.20983, Z=3.765, p<0.0001665, RF: Median=1.20967, Z=3.454, p<0.0005516).

This overestimation means that after trial 5 the participants IMI is longer than the period of the metronome. This can explain in turn the positive retraction of the within-block SMA curves after block-trial 5, seen in Figure 3B. As the maximum negative SMA occurs in trial 4, then from trial 5 onwards, as the participants start moving with an IMI larger than metronome s period, their movements start getting closer to the metronome’s stimulus, becoming progressively less negative. The fact that after block-trial 5 the IMI remains consistently larger than the metronome, with an almost constant (very slow), overestimation and that the SMA retains a monotonic positive slope without corrective deflections, suggests that the second part of the block is primarily concerned with maintaining a constant IMI rather than maintaining SMA at a specific level.

#### Across-block IMI

The across-block IMI was calculated by first computing for each subject and limb the average IMI within each block and is shown in Figure 4C. In order to focus on the last trials of the blocks where the period overestimation is evident, only trials 7,8,9 and 10 were included in the computation of each block average. Then the single-subject, across-block curves were averaged across subjects for each limb and provided the overall mean IMI, across-block curves with the corresponding standard error. As it can be clearly seen in overall the IMI stays higher than 1.2sec for the entire experiment. In order to test if there is a significant reduction of this overestimation in the later part of the experiment, a Wilcoxon signed rank test was performed between the average across-block IMI in the first half (run 1) and in second half (run 2) for each limb. The significance level was adjusted for the 4 different tests, one for each limb, to 0.0125. No significant differences were found between the two halves of the experiment for the overestimated IMI. Namely, (LH: Z=-2.229, p<0.0258385, RH: Z=2.087, p<0.0369246, LF: Z=1.129, p<0.2587892, RF: Z=-0.213, p<0.8312920).

In addition, we performed in each of the halves a Wilcoxon test per limb in order to verify that in both halves the overestimated IMI was significantly higher than the metronome’s period. The significance threshold was adjusted for the 8 different tests (4 limbs, 2 runs) to 0.0063. For all limbs and in both halves the IMI was significantly higher that the metronome’s period of 1.2 sec. Namely for the first half (LH: Median=1.20795, Z=4.371, p<0.0000124, RH: Median=1.20928, Z=5.032, p<0.0000005, LF: Median=1.21637, Z=4.788, p<0.0000017, RF: Median=1.20812, Z=3.862, p<0.0001126). And for the second half (LH: Median=1.21318, Z=4.446, p<0.0000088, RH: Median=1.20593, Z=3.221, p<0.0012753, LF: Median=1.21329, Z=4.894, p<0.0000010, RF: Median=1.20969, Z=3.238, p<0.0012044).

The fact that across blocks there is no gradual reduction (or very slow so that is not captured by the statistics) of the overestimation towards the actual period of the metronome, hints towards a systematic type of bias rather than a bias reflecting unfamiliarity with the given duration. If this were the case then the brain would need only a few blocks of exposure to the constant metronome’s period to correctly capture it and reduce the overestimation. As there is no reduction it seems that this is a persistent bias.

Overestimation of the metronome’s period by the IMI has already been observed in previous research. Michon (Michon & van der Valk, 1967) showed that during steady-state sensorimotor synchronization, a step increase in the metronome period, causes an IMI overshoot in the following trials which relatively quickly, within a few trials, converges to the new period. Thaut and colleagues (Thaut et al., 1998) did a similar experiment by parameterizing the percentage of the period step change w.r.t to the metronome period before the change. They showed that in all examined levels of step change there was an overshoot of the IMI and then gradual decline towards the new period. For small percentage changes, the upward overshoot phase period lasted for 6-7 trials while for larger percentages it was faster. Repp (Repp, 2001) also investigated the adaptation of IMI after step changes of the metronome’s period and confirmed the exact same pattern, with one of the differences being that for small step changes, the IMI curve increased smoothly without a large overshoot to a plateau of overestimated period for as long as 10 trials. This pattern resembles the IMI curve observed here. The above mentioned studies involved a change of the metronome period during steady-state synchronization. In our study the adaptation starts from rest and it could be the case that under such conditions, it is as easy for the brain to adapt to a new rhythm as easy it is to adapt to a slight increase of an ongoing steady state rhythm. This speculative hypothesis would explain the existence or absence of overshoot and the long-sustained overestimation in IMI.

#### IMI overall differences between limbs

The overall mean IMI was computed for each limb in order to examine the differences between limbs. This was performed by first computing for each participant the mean IMI across the entire experiment for each limb. Then for each limb the overall mean IMI was computed across subjects together with the standard error of the mean. These statistics are shown in Figure 4D. The mean IMI values for individual subjects were used for performing Wilcoxon signed-rank tests between all combinations of limb pairs. The Wilcoxon test was chosen because, similarly to SMA, the distribution of the mean IMI for all subjects was found to not resemble a normal distribution for any limb, as indicated by the Chi-Square test for normality.

According to the Wilcoxon test, no significant differences were found between Left and Right Hands (LH-RH: Z=1.228, p<0.2196217). Between Left and Right Feet there was a significant difference, even when the significance level (0.05) was Bonferroni corrected for the 6 different tests, leading to a significance level 0.0083 (LF-RF: Z=2.838, p<0.0045452).

In the comparisons between Hands and Feet the situation was very different from the SMA case. Here 3 tests were found non-significant (LH-LF: Z=-2.602, p<0.0092801, RH-RF: Z=-1.043, p<0.2967755, LH-RF: Z=0.210, p<0.8336181) and only one significant between Right hand and Left foot (RH-LF: Z=-4.244, p<0.0000220) which are the two cases with the biggest separation in Figure 4D.

These overall statistics show that for the IMI there is no clear pattern that distinguishes hands and feet. This of course is also obvious from the within-block average curves in Fig. 4B, where there is no obvious difference between any of the limbs. And from the across-blocks average curves in Fig.4C, where it can be seen that in the first half of the experiment there seems to be some difference between hands and feet which then disappears in the second half of the experiment.

### VARIABILITY of IMI

In order to study the variability of IMI we first computed in each trial of the experiment the standard deviation of IMI across subjects for each limb. Figure 4E presents the standard deviation curves for all limbs across all the trials of the experiment. From this plot the most obvious phenomenon is almost the entire length of the experiment the hands (red and green) have lower variability than feet (blue and grey).

#### Within-block behavior of IMI Standard Deviation

The within-block curve of standard deviation was computed similarly as for the standard deviation of SMA. IMI is defined for only 9 trials in the block, trials 2 to 10. The average within-block standard deviation curves for all limbs are shown in Figure 4F. All limbs have a decreasing convex curve.

The first IMI in trial 2 in the block has the largest standard deviation. Then the second IMI in trial 3 gets smaller and the third IMI in trial 4 decreases further to a level around which settles the standard deviation of the following IMIs, in trials 5 to 10.

This pattern of within-block IMI standard deviation behaves in the opposite way of the standard deviation of the SMA (comparing Figures 3F and 4F). At the beginning of the block the variability of SMA is low while the variability of IMI large. As the block progresses, the SMA becomes gradually more variable and the IMI less variable. Then around trials 4 to 5, the SMA standard deviation converges to a maximum plateau while IMI settles around the same trials to a minimum plateau. This behavior could be intuitively described as the brain putting more emphasis on being accurate on SMA at the very beginning of the block and then gradually up to trials 4 to 5 switching mode, thereafter putting more emphasis on being accurate on IMI.

Hands have consistently smaller IMI standard deviation than feet in the entire block without any obvious convergent or divergent behavior within the block.

#### Across-blocs behavior of IMI Standard Deviation

The across-block IMI curve of standard deviation was computed similarly to that for SMA. The curves for all limbs are shown in Figure 4G. There are two clear opposite patterns for hands and feet. For the hands the standard deviation of IMI asynchrony starts with low values in the first blocks and progressively increases to higher values. The opposite occurs for feet for which IMI has high standard deviation at the beginning of the experiment and then it progressively has less and less variability. All limbs seem to converge to a similar standard deviation level at the end of the experiment. The patterns for left and right hands are identical and the same holds for feet.

By directly comparing Figures 3G and 4G it can be seen that this pattern is strikingly similar to the across-block behavior of the standard deviation of SMA.

So while across blocks the variability of SMA and IMI follow a similar pattern, within-block they follow a completely opposing pattern. This is a manifestation of different stochastic sources contributing to the within-block and across-blocks variability.

#### IMI Standard Deviation difference between Hands and Feet

The overall standard deviation of IMI was computed for each limb in order to examine the differences between limbs. This was performed in a similar way as for the SMA and the results are shown for each limb in Figure 4H. The values for all subjects were used for performing Wilcoxon signed-rank tests between all combinations of limb pairs similar to the tests for the SMA.

According to the Wilcoxon test, no significant differences were found between Left and Right Hands (LH-RH: Z=1.281, p<0.2002221). In contrast, significant difference was found between Left and Right Feet (LF-RF: Z=3.255, p<0.0011352).

All comparisons between Hands and Feet were found significant (LH-LF: Z=-5.710, p<0.00000001, RH-RF: Z=-5.677, p<0.00000001, LH-RF: Z=-4.566, p<0.0000050, RH-LF: Z=-6.235, p<0.00000001) These results were expected, as in both the within- and between-blocks curves a consistent difference between hands and feet was seen.

### Summary of results for IMI

Three main patterns were observed in the IMI:

1. Within block, IMI has an increasing convex form, starting from a low value (justified by the movement in the first block trial being a response and in the second block trial an anticipation) and it gradually increases up to the metronome s period by trial 5. This slow increase does not become faster in the later parts of the experiment but remains slow. Thereafter the IMI settles at a value slightly but significantly higher value than the metronome’s period. This is a phenomenon that occurs in both halves of the experiment. This pattern is very similar for all limbs.
2. Across blocks, in the long range of tens of minutes, the IMI remains higher than the metronome’s period and no convergence towards it is observed. This is evidence that this overestimation corresponds to a bias and not to an unfamiliarity with the given metronome’s period which can be learned and adapted to. This pattern is very similar for all limbs.
3. Within and across blocks, in overall, no consistent average IMI differences were found between limbs.

The analysis of the standard deviation of IMI revealed 3 main interesting phenomena:

1. Within block, the first IMI in trial 2 in the block has the largest standard deviation. Then the second IMI in trial 3 gets smaller and the third IMI in trial 4 decreases further to a level around which settles the standard deviation of the following IMIs in trials 5 to 10. This is the exact opposite behavior of the standard deviation of the SMA, which is smallest at the beginning of the block and progressively increases up to trial 5 where it settles at a maximum plateau.
2. Across blocks now, for the hands the standard deviation of IMI asynchrony starts with low values in the first blocks and progressively increases to higher values. The opposite occurs for feet for which IMI has high standard deviation at the beginning of the experiment and then it progressively has less and less variability. This pattern is similar to the across-block behavior of the standard deviation of SMA.
3. Feet have on overall higher IMI variability than hands.

## THE NEGATIVE SLOPE OF THE EVOLUTION OF SMA ACROSS BLOCKS

The most striking phenomenon in the results of the current work is the near-linear negative gradient of the SMA across blocks for all limbs.

As it has already been described, the presentation of the blocks for a specific limb was not contiguous but between every two successive blocks there was a number of interleaved blocks or other limbs or of resting. For example consider the case of Left Hand blocks (green rectangles in the sequence of Figure 1B). Between the first and second block of Left Hand there were 6 blocks of other types. Between the second and third block of Left Hand there were 5 blocks of other types. Between third and fourth block of Left Hand there was 1 block of other type. With such variability between successive blocks of the same limb there are three possibilities of how SMA would be expected to behave.

1. The SMA would be reset and randomized every time there is change of limb in the block sequence. This reset and randomization would occur around a baseline level of negative SMA, constant across the experiment and characteristic for each limb, with similar baseline values for the two hands and the two feet respectively. This behavior would correspond to a sensorimotor synchronization system without any within- and between-limb memory. The constant baseline level of the negative SMA would represent the average anticipatory motor lead, superimposed on the kinematic/kinesthetic characteristics of the given limb.
2. The SMA of the current block would serve as the baseline for the next block, irrespective of the limb. Based on this baseline the participant would become more anticipatory in the following block, leading to an increase in the SMA (more negative). With a fixed such rate of increasing asynchrony it would be expected to have a monotonically increasing, near-linear, negative mean asynchrony until a final target steady state level has been reached. This would correspond to a brain mechanism in which sensorimotor synchronization is a common process for all limbs underlined by short-term memory transfer between them. Under such mechanism there is a clear prediction that the SMA precession across blocks is proportional to their number, irrespective of limb type. So the SMA increase between two successive blocks of Left Hand interleaved by 6 blocks of other limbs should be double that of the increase across 3 interleaved blocks and triple that of the increase across 2 interleaved blocks.
3. Similarly to above, the SMA of the current block would serve as the baseline for the next block, but only for the same limb. Based on this baseline the participant would become more anticipatory in the next block of only the same limb. Again, it would be expected to have a monotonically increasing, near-linear negative mean asynchrony across blocks until a final target steady state level has been reached. But in this limb-specific case the SMA precession across blocks would not be affected by any interleaved blocks of other limbs. This means that this precession between two successive blocks of Left Hand would be the same whether there were 2, 5 or 8 interleaved blocks of other limbs. This behavior would correspond to a brain mechanism in which sensorimotor synchronization is a limb-specific process with short-term memory spanning several blocks. It is important to highlight that in such scenario each limb has each own sensorimotor short-term memory.

Out of these three possible mechanisms of sensory-motor synchronisation, it is clearly evident that the first mechanism cannot capture the monotonic evolution of the SMA curves across blocks in Figure 3C. As there is no SMA reset or randomization in each block, this means that there is some form of memory which underlies the gradual descent of SMA across blocks. The other two possible mechanisms have very distinct memory characteristics. Either a common memory mechanism for all limbs in which SMA gradual precession occurs in a contiguous way, so that all limbs affect all limbs. Or a limb-specific memory which updates the SMA precession only in blocks of the same limb.

In order to disentangle which of these two types of memory underlie the negative gradient of SMA across blocks, an one-way analysis of its variance was performed. The independent variable in this analysis was the distance (in blocks) between successive blocks of the same limb.

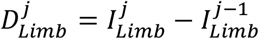

In the above equation there are two types of block indices. The index *j*, termed here as “limb-block” index, refers to the index of a block only within the sequence of the blocks of the same limb. So, for a specific limb the limb-block index *j* takes values between 1 and 16. The second type of index *I*, termed here “total-block” index, refers to the index of a block in the entire experiment sequence of blocks including all limbs and the resting blocks. So total-block index *I* takes values between 1 and 64. For example for the Right Hand the first block, limb-block index *j* = 1, occurs in the fourth experimental block and has a total-block index *I* = 4. The second Right Hand block, *j* = 2, occurs in the sixth experimental block and has a total-block index *I* = 6. So the distance between these two successive blocks of Right Hand is 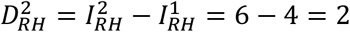 blocks. In this fashion the block distance between every two successive blocks of every limb, *Limb* = {*LH, RH, LF, RF*}, were computed.

The dependent variable for the analysis of variance was the difference of SMA between successive blocks of the same limb.

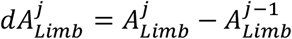

where *A* is the average SMA in a block and *j* is the limb-block index of a block for a specific limb, as described just above.

Regarding the independent variable, 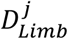, a very important aspect is that the experimental design of this task in the Human Connectome Project was not developed to be balanced with respect to this specific parameter. In order to demonstrate this issue, Figure 6 shows for each limb the number of occurrences of different distances between successive blocks of the same limb. For example for Left Hand, there are 4 instances when the distance between two successive left-hand blocks was 2 blocks, 6 instances when the distance was 6 blocks, and 4 instances when the distance was 7 blocks. There are no instances when the distance was 3,4,5, or 8 blocks. Different patterns occur for different limbs.

**Figure 6.**
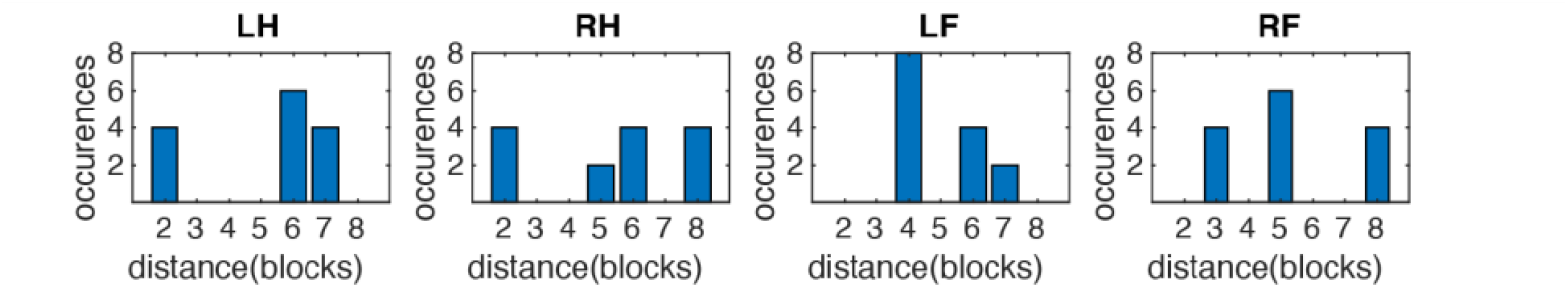
Number of interleaved blocks of other limbs between successive blocks of the same limb. (LH,RH: Left/Right Hand, LF,RF: Left/Right Foot). Each limb has a unique pattern of unique distances (in blocks) between successive blocks of the same limb. On Y axes is depicted how many times a unique distance occurs across the experiment.

Here it must be noted that devising a similar experimental design with balanced occurrence of distances of interleaved blocks between successive blocks of the same limb should be considered in some aspects impractical. Even if only the distances of 1,2 and 3 blocks would be considered in an experiment involving all 4 limbs, this would require a very long sequence of blocks in order to provide the balanced experimental structure. For example such a sequence could be constructed based on graph theory as a de Bruijn pseudo-random sequence (de Bruijn, 1946) and would consist of 256 blocks. Given that each block is about 16 seconds long, this sequence would require about 64 hours of pure experiment time per participant, something impractical. So the current experimental design, although unbalanced, still offers the opportunity to study resiably how sensory-motor memory of the SMA behaves when other limb movements are interleaved.

Despite this unbalanced design, the wide range of distances that occur for each limb offers the possibility to investigate whether the increase of the SMA is proportional to block distance or not. In the case that it would proportionally increase with distance, this would be evidence that the interleaved blocks have an effect in the gradient of the asynchrony and that the sensory-motor memory mechanism is common between limbs. In the case the asynchrony would not increase proportionally to distance, it would mean that the sensory-motor memory is limb-specific and all interleaved other limb movements have no reinforcing effect on it.

The only major confound that could affect such analysis is when the number of occurrences of the different distances monotonically increases or decreases with distance. Investigating Figure 6, it can be seen that this only occurs for the Left Foot case, where there are 8 occurences of the distance of 3 blocks, 4 occurrences of distance of 6 blocks and 2 occurences of distance of 7 blocks. This monotonic decrease of occurences could potentially introduce a confound and result in a slope of asynchrony increase w.r.t to distance, just due to the different levels of bias in each distance. So in case of a significant slope for this limb, special consideration should be taken due to the methodological limitations. For the other limbs there is no such issue, as the number of occurrences does not increase or decrease with distance but varies in comparable levels(see figure 4A).

The first obvious choice for the analysis of variance was a first-order fixed-effect ANOVA but before employing it the first step was to examine whether the dependent variable 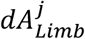 follows a normal distribution. This was performed by applying a Chi-Squared Goodness of Fit Test (Balakrishnan et al., 2013)with the null hypothesis being that the distribution of the difference of SMA between successive blocks of the same limb, 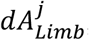, follows a normal distribution. For each limb all these asynchrony difference values across blocks were collected for all subjects and the Chi-Squared Goodness of Fit Test was performed on them. Four such tests were performed in total, one per limb. The significance level 0.05 was adjusted accordingly through Bonferroni correction to *a*_*c*_ =0.0125. In all four tests the null hypothesis, that 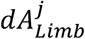 followed a normal distribution, was rejected at the significance level *a*_*c*_.

Namely:

Left Hand: *χ*^2^(5, *N* = 854) = 30.70, p = 1.4*10^−11^, *H*_0_ rejected

Right Hand: *χ*^2^(5, *N* = 854) = 36.12, p = 2.55*10^−11^, *H*_0_ rejected

Left Foot: *χ*^2^(5, *N* = 854) = 71.17, p = 3.10*10^−18^, *H*_0_ rejected

Right Foot: *χ*^2^(5, *N* = 854) = 94.34, p = 9.01*10^−17^, *H*_0_ rejected

This very significant deviation of 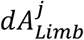 from normal distribution is evident in Figure 7, where it is shown that its distribution is leptokurtic (peakier with heavier tails) with much higher kurtosis than the normal distribution.

**Figure 7.**
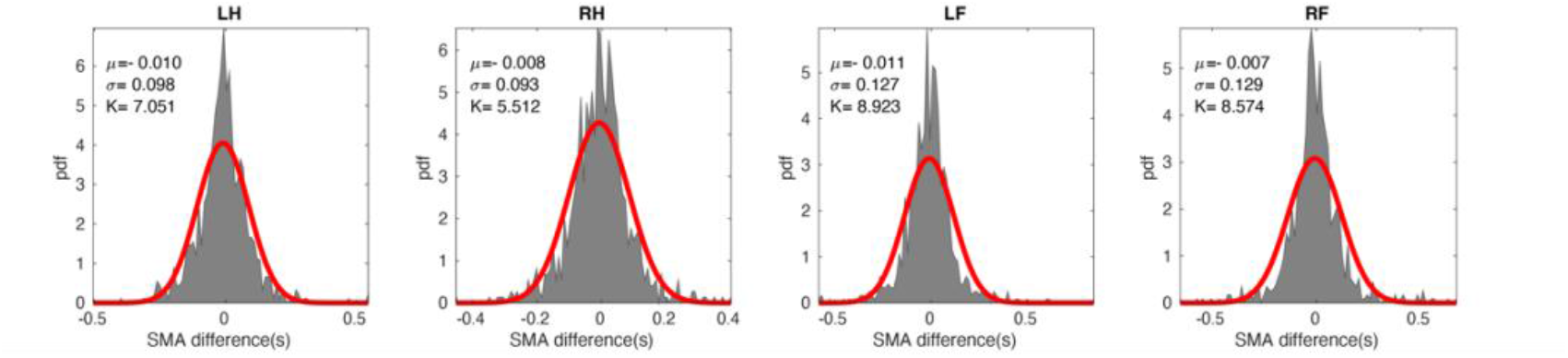
Distribution of SMA differences across successive blocks of the same limb. (LH,RH: Left/Right Hand, LF,RF: Left/Right Foot). The red line is a fitted normal distribution. The deviations of the data from the normal distribution are obvious. This is also captured by the kurtosis K of the data histograms which is much higher than 3, that of the normal distribution.

We then as a sanity check performed for each limb a Wilcoxon test in order to verify that the median of the SMA difference 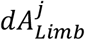 was significantly different from zero, as this is obvious from the monotonic increase (more negative) of SMA, 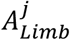. The Wilcoxon test was selected because it is a non-parametric test and does not assume normality of the data. Four such tests were performed, one for each limb and the default significance level was again Bonferroni-corrected accordingly to *a*_*c*_ =0.0125. The tests confirmed for each limb the median of 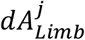 is significantly different than zero and negative as expected. Namely:

Left Hand: *M=*-0.008s *H=*1 *Z=*-3.237 p<0.0012

Right Hand: *M=*-0.005s *H=*1 *Z=*-2.522 p<0.0116

Left Foot: *M=*-0.009s *H=*1 *Z=*-3.243 p<0.0011

Right Foot: *M=*-0.008s *H=*1 *Z=*-2.552 p< 0.0106

For the analysis of variance, due to the strong deviation of the dependent variable from the normal distribution, it was decided instead of ANOVA to use the non-parametric Kruskal-Wallis test by ranks(Kruskal & Wallis, 1952). Four tests were performed, one for each limb. In each such test the number of groups was equal to the number of unique block distances between successive blocks of the given limb. For Left Hand there were 3 groups of block distance, namely [2, 6,7] blocks. For Right Hand there were 4 groups of block distance, namely [2, 5, 6, 8] blocks. For Left Foot 3 groups of [4, 6, 7] blocks and for Right Foot 3 groups of [3, 5, 8] blocks.

For each limp the null hypothesis *H*_0_ of the Kruskal-Wallis test was that for each group of block distance the SMA difference comes from the same distribution. This would mean that the SMA difference between successive blocks of the same limb is not affected by the number of interleaved blocks of other limbs.

The alternative hypothesis *H*_*a*_ was that the SMA difference comes from different distributions for different blocks distances between successive blocks of the same limb. This would be the case, if the SMA difference would increase proportionally to the number of the interleaved other-limb blocks so that it would be bigger for longer block distances.

The Kruskal-Wallis tests showed that there was no statistically significant difference of 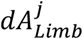 for different distances. Namely

Left Hand: H(2)=0.974, p=0.614, *H*_0_ NOT rejected

Right Hand: H(3)=1.384, p=0.709, *H*_0_ NOT rejected

Left Foot: H(2)=1.640, p=0.440, *H*_0_ NOT rejected

Right Foot: H(2)=3.018, p=0.221, *H*_0_ NOT rejected

The lack of effect of the block distance on 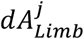 is evident in Figure 8 A to D, where the boxplots show the median and 25% and 75% percentiles at each block distance. If the SMA would grow with block distance then 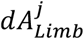 should be progressively more negative with increasing block-distance. As it is evident from the plots that there is no such tendency. It is characteristic that in post-hoc tests of 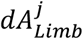 between each possible pair of block-distances, no pair was found with statistically significant difference for any limb. Not even pairs between the extrema of distances, i.e. between distances of 2 and 7 blocks for left hand.

**Figure 8.**
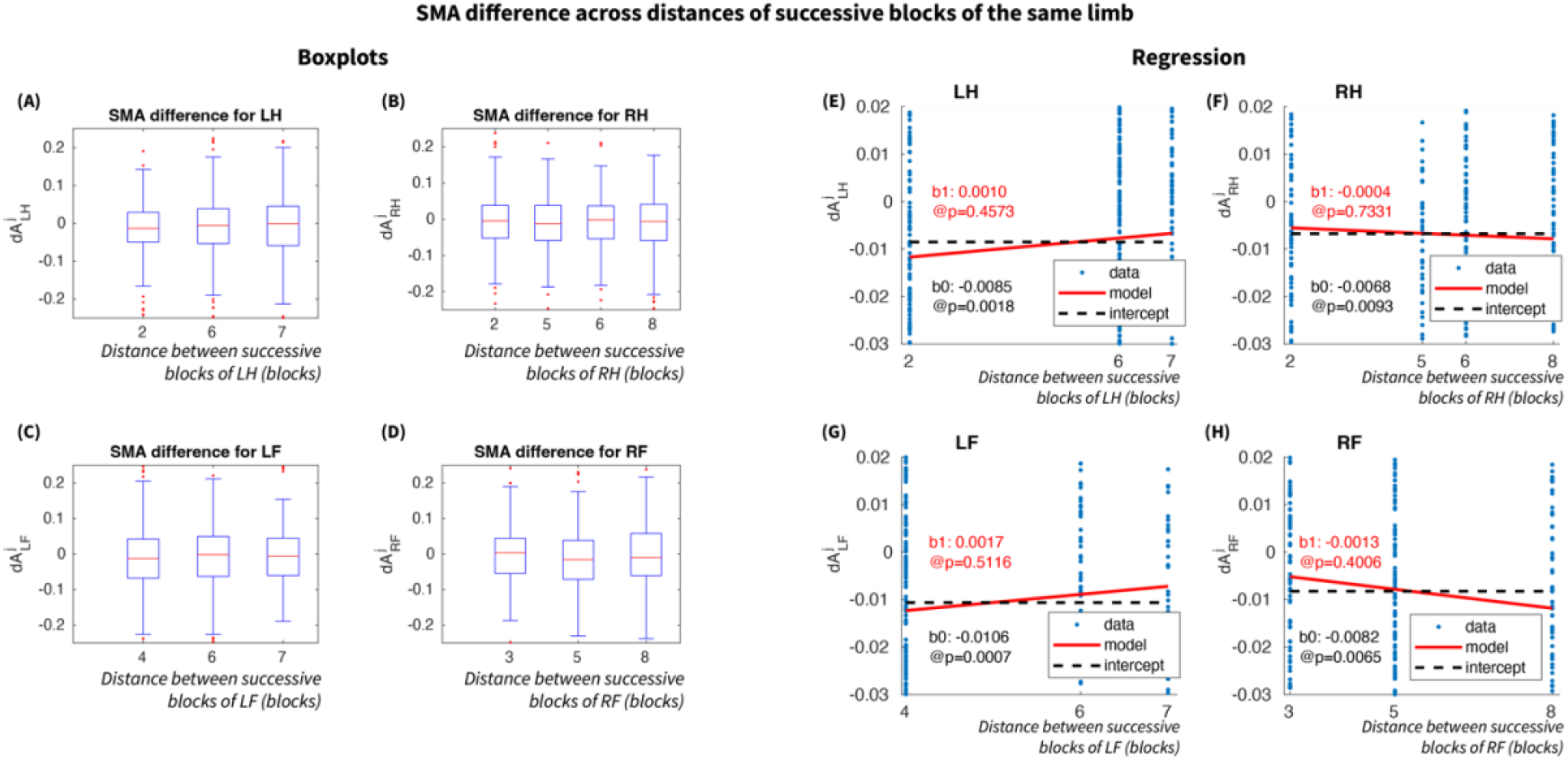
SMA difference across distances of successive blocks of the same limb. (LH,RH: Left/Right Hand, LF,RF: Left/Right Foot) (A)-(D): Boxplots of SMA differences across different distances of same limb successive blocks. No significant differences were found across any distance combinations. This means that the number of interleaved blocks of other limbs does not affect the SMA difference between successive blocks of the same limb. The mean SMA difference is significantly negative for all distances with an overall mean of −7 msec. (E)-(H): Regression was performed between SMA difference and distance in order to verify that there is no significant slope across distances. The estimated slopes with their corresponding p-values are printed in red on the plots. For all limbs the slope was not significantly different than 0. The intercept is displayed with dashed black lines and its value and corresponding p-value are printed in black on the plots. All intercepts were significantly negative. These analyses showed that the number of interleaved blocks of other limbs does not affect the gradient of SMA increase(more negative) across blocks of the same limb. This hints to SMA being a limb-specific process, involving short-term memory.

As a sanity check of this observation linear models of the form *dA*_*Limb*_(*j*) = *b*_0_ + *b*_0_ · *D*_*Limb*_ (*j*) were fitted, where *b*_0_ is the intercept and *b*_0_ is the slope coefficient. It was expected that the intercept *b*_0_ would be significantly negative as it captures the average difference of 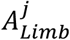 across blocks, which is consistently negative. It was also expected that the slope *b*_0_would not be significantly different than zero as no gradient of 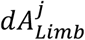 was seen in the boxplots with increasing block distance. Both these expectations were confirmed. The fitted models for each limb are presented in Figures 8 E to H. On these plots one can see that all intercepts (dotted black lines) are significantly negative for all limbs, while the slope coefficients are not significantly different from zero (red lines). The intercepts values were found to be LH: −0.0085, RH: −0.0068, LF: −0.0106, RF: −0.0082 sec.

The fact that there is no effect of block-distance on 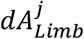 is a novel finding with direct implications on the characteristics of the memory mechanism employed in such a simple sensorimotor task. The fact that 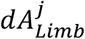 remains the same irrespective of the number of interleaved blocks of other limbs is strong evidence that the memory is limb-specific and is not affected by what happens in the other limbs. And this short-term memory can persist in the order of minutes as it is demonstrated by the longest block-distances in this experiment of 7 or 8 blocks (each block had a duration of 15 seconds).

## THE WITHIN-BLOCK INSTANTIATION OF THE SMA EVOLUTION ACROSS BLOCKS

Having established that the negative SMA gradient across blocks is limb-specific, the next important question is whether this gradient is instantiated in every block-trial within a block. The SMA gradient across blocks, shown in Figure 3C, represents the average of each block. However, it is still not known whether this across-block gradient is the same when instead of the mean of each block, we examine each block-trial individually (trials 2 to 10, as trial 1 in each block movement is always a reaction). One possibility would be that the across-blocks gradient is different for each trial in the block. For example the gradient could be steepest for trial 2 and progressively flatter for the remaining trials in the block, reflecting a more dominant role of the early part of the block. Another case could be that the gradient in trial 2 starts from a small value and progressively increases in each trial, reflecting a more distributed role of each trial.

Do resolve this gradient of the SMA across blocks was examined in each block-trial position. The SMA was split in 10 bins, one for each block-trial. Then in each bin the SMA was averaged in each block across subjects. This provided for each block-trial a time-series with 16 average SMA values across the 16 blocks. These timeseries for each of the 10 block-trials are shown for each limb in Figures 9 A to D for each limb. The most obvious phenomenon on these plots is that for trial 1 the SMA is highly positive and remains flat, around the same level across blocks. This was expected as the movement in the first trial of each block is a reaction and there is no anticipatory element involved. The curves for all the other trials have a clear negative slope becoming more negative with elapsed blocks, similar to the behavior of the block average. Also the offsets of these curves reflect the already described “hook” shape of the within-block SMA behavior. After the highly positive flat curve for trial 1, the curve offset reaches a minimum for trial 4 and then increases again up to trial 10.

**Figure 9.**
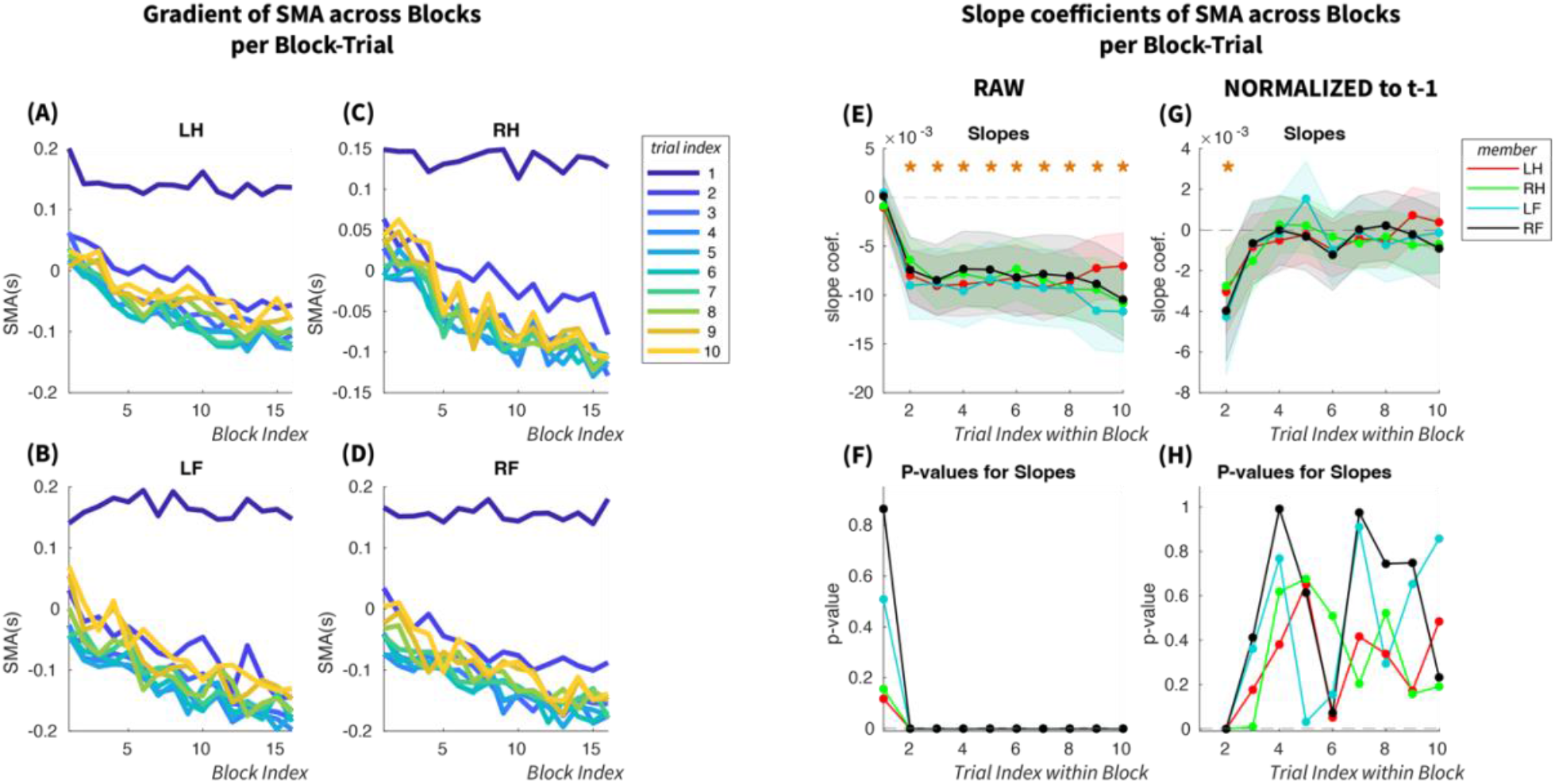
Across-Blocks SMA Gradient per Block-trial. (LH,RH: Left/Right Hand, LF,RF: Left/Right Foot) (A)-(D): Across-Blocks gradient of SMA at each Block-Trial. SMA in Block-trial 1 has a flat slope across blocks because it is a reaction time without anticipatory effects. For all other Block-Trials the gradient across blocks is clearly negative. There is no obvious difference between these gradients. (E) Slope coefficients of a linear model fitted on the gradient curve for each Block-trial in plots (A)-(D). The slope for Block-Trial 1 was not significantly different than 0. For all other Block-Trials the slopes were significantly different than 0 (signified by yellow stars), around a steady level without any monotonic decrease across Block-Trials. (F) The corresponding p-values of the slopes that confirm the significance for Block-Trials 2 to 10. (G) SMA normalized to the previous Block-Trial. In each Block-Trial the SMA gradient across blocks is normalized by subtracting the SMA gradient of the previous Block-Trial. This shows how much of the overall gradient remains after the gradient of the previous Block-Trial is removed. The results show that only in Block-Trial 2 there is a significant gradient. From Block-Trial 3 onwards the remaining slope is not significantly different than zero. This shows that the SMA gradient across blocks is mostly explained by the gradient of Block-Trial 2 and the remaining Block-Trials do not contribute to it. (H) Corresponding p-values for the normalized slopes. Only in Block-Trial 2 the p-value is significant.

Regarding the slopes of the curves, they seem on the plots to be comparable for trials 2 to 10. For trial 2 the similarity with the other trials is not so obvious due to the curve offset. In order to quantify this analysis, a linear model was fitted on each curve from which the slope coefficient was used to quantify the gradient of the curve for each trial. Each model was of the form:

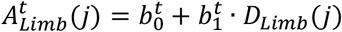

where integer *t* represents the within-block trial index, *t* ∈ [0,00], 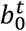 is the model intercept and 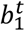 the model slope coefficient.

The slope coefficients are presented in Figures 9E, with their 95% confidence intervals. Apart from trial 1 where the slope is close to 0, for all the other trials the slope is significantly negative and remains around the same level without any monotonic increase or decrease across blocks. These slope coefficients were highly significant. As there are 10 trials and 4 limbs there are 40 slope coefficients and corresponding p-values and thus the significance level has been adjusted with Bonferonni correction to *p*_*threshold*_ = 0.05*/*40 = 0.00125. For all the slope coefficients for trials 2 to 10 and all limbs the p-value was smaller than 1.217*10^−6^. This is depicted in Figure 9F, where it can also be seen that for trial 1 the p-values are very large, as the slope is near zero for the first block-trial.

In order to examine quantitatively whether the slope remains the same across trials or becomes from trial to trial steeper of flatter, we fitted linear models to the SMA difference between successive trials. The idea behind this is that from each trial we remove the slope of the previous trial and then we test whether the remaining slope is still significantly different than zero. So the dependent variable here is defined as

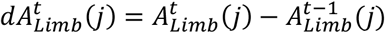

where the trial index here can take values *t* ∈ [2,00]. For each limb there were 9 such time-series, as for trial 1 this difference is not defined, giving for all limbs 36 time-series. Each such time-series had 16 values, one for each block. On each of these time-series was fitted a linear model of the form:

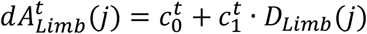

The slope coefficients 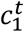 of these models are plotted on Figure 9G, with their corresponding 75% confidence intervals. The p-values of these coefficients are plotted on Figure 9H. From these plots it is obvious that only in trial 2 there is a significant negative slope. All other trials had coefficients not significantly different from 0. For assessment of significance, the p-value threshold of 0.05 was Bonferroni corrected for the 36 different fitted models as *p*_*threshold*_ = 0.00038. For trial number 2 the p-values of the slope coefficients were LH: p=0.00016, RH: p=0.00017, LF: p=0.00011, RF: p=0.00004, all smaller than the significance threshold. For all other trials the p-values of the slope coefficients were at least one order of magnitude larger than the significance threshold.

The above results show that the near-linear negative gradient of the SMA across blocks is only instantiated only in trial 2 of each block, the first trial in which the movement is anticipatory. All subsequent trials, 3 to 10, in each block do not contribute anything the evolution of the SMA to more negative, anticipatory values across blocks. This finding is novel and non-trivial. It means that after the reactive movement in trial 1, the brain engages its anticipatory mechanism, responsible for the long-term negative asynchrony gradient, only in trial 2 and after that it disengages from it. Intriguingly this directly implies that in all subsequent trials, 3 to 10, the brain switches to a different task. And the obvious candidate for this task is the alignment of the Inter-Movement Interval with the metronome’s period.

### Summary of Results from the investigation of the negative slope of SMA across blocks

There were two novel and significant findings in the investigation of the negative gradient of the SMA across blocks.

1. The gradient of SMA across two successive blocks of the same limb does not depend on the number of interleaved blocks of other limb movements. It appears to have an average gradient of about −8 msec whether there are few or many interleaved blocks. More importantly this is direct evidence that there is limb-specific-short term sensorimotor memory which persists in the order of tens of seconds.
2. The negative gradient of SMA across blocks is instantiated only in the first anticipatory movement (in trial 2 of each block). The following block trials (3 to 10) have no contribution to the negative gradient of the SMA. This is direct evidence that in each block the brain engages for only one trial in the task of adjusting its anticipatory movement phase with respect to the metronome and that in the rest of the trials it disengages from it and engages in a secondary task. This secondary task appears to be the alignment of the Inter-Movement Interval with the metronome’s period.

## DISCUSSION

There are three main overarching contributions of this work. The first contribution is the existence of three distinct temporal scales of sensorimotor synchronization with distinct signatures. A long-range, across-blocks monotonic negative gradient of SMA to more anticipatory movement, which prevails for tens of minutes, a very consistent “hook”-shaped pattern of SMA within each block, in the range of seconds, and a constant SMA difference across time between feet and hands. The second contribution is the demonstration that the across-blocks, monotonic, negative gradient of SMA to more anticipatory movement is instantiated only in the first anticipatory trial of each block (trial 2) and the rest of the subsequent block trials have no contribution to this SMA gradient. The results here suggest that in these trials the “hook”-like asynchrony shape of SMA is driven by the alignment of the IMI to the metronome’s period. The third contribution of this work is that this negative SMA gradient is limb-specific and is not affected by the interleaved blocks of other limbs.

These findings have very important implications about the functional principles employed by the brain during sensorimotor synchronization to external rhythmic stimuli. The main assumption of most influential models of steady-state sensorimotor synchronization is that SMA and IMI corrections are using information only from the immediate previous one or two trials, with an autoregressive mechanism of very limited memory(Jacoby & Repp, 2012; Repp, 2005). These models fail to capture the behavior of SMA and IMI of the current study during the “tuning-in” phase of sensorimotor in multiple aspects.

Here, the SMA has a strong short-term memory component with a long span across blocks, in the order of minutes, which drives the negative slope across-blocks. More importantly this long memory component is limb-specific and when blocks from other limbs are interleaved, the asynchrony is not affected but it rather seems to be put on hold until the next block of the same limb. In order to capture this behavior new types of models need to be developed, which will include a non-linear component that will capture the hold-and-retrieve, limb-specific memory of the asynchrony. No such short-term memory pattern was observed for IMI.

In addition to the long-term memory component of SMA, there seems to be a clear distinction between SMA (phase) and IMI (period) in the priority by which they are tuned to the metronome in each block. When a block starts, the participant waits for the first stimulus in order to start moving so the movement there is a reaction rather than an aniticipation. Then the anticipatory mechanism is deployed and the participant does not wait for the second stimulus in the block but knowing well the period of the metronome it moves before the second trial occurs. For this first anticipatory movement the brain recalls from memory the stored information of the SMA in the previous block of the same limb and advances it to an even more anticipatory, more negative, level. This is the only trial where the task of aligning the SMA to a mental target occurs. Then the brain switches task and between trials 3-5 its task becomes to align the IMI to the metronome’s period. Once this is achieved by trial 5 then for the rest of the block the brain just maintains this IMI stable. So from this mechanistic description in each block the primary task of aligning the asynchrony occupies the brain for 1 trial while the secondary task of aligning the IMI occupies 8 trials. This inbalance in spent resources by the brain in the two different tasks might be revealing about the reason behind the across-block negative slope of SMA. Although the brain has an inherent automatic tendency to become more anticipatory and thus have a more negative asynchrony, once it locks to a specific phase w.r.t the stimulus it switches its focus to aligning its IMI to the period of the metronome and maintaining it. So although the period alignment is the secondary task in terms of sequence of instantiation in each block it seems to be the more cognitively important of the two as the brain allocates the vast majority of resources to it.

Another piece of evidence supporting the switching of focus between SMA and IMI within a block is the opposite behavior of their within-block standard deviation curves. The SMA curve starts with low variability and progressively increases until it reaches a maximum plateau while the IMI starts with high variability and progressively decreases down to a minimum plateau in trial 4. This opposing behavior appears to fit well with the proposal that at the beginning of each block the brain has as primary task the SMA and that is why it is less variable but it quickly switches to the task of aligning IMI to metronome’s rhythm so IMI becomes the more accurate task.

The distinction of SMA and IMI alignment as two separate processes is not a completely new idea. It has already been proposed in previous research, given some evidence, that these two tasks might employ different brain networks(Middleton & Strick, 2000; Repp, 2000, 2001; Repp & Keller, 2004). The results of the current study contribute substantial evidence in favor of this distinction. What is trully new is the evidence that the brain shows first a strong instinctive tendency to become anticipatory, by adjusting SMA, but then it quickly changes focus to period alignment and remains engaged to it. It seems that asynchrony alignment is an automatic process that utilizes short-memory while period alignement is a more conscious process driven by the task at hand and does not utilize memory, if it does not have to. Evidence for this latter part comes from the fact that within-block the convergence of the IMI to a plateau near the metronome’s period does not become faster in the second part of the experiment, as compared to the first. In both halves the IMI curves look identical, reaching the vicinity of the metronome’s period in trial 5, while one would expect that after long exposure to the constant rhythm of the metronome, the participants in the second half of the experiment should be able to converge faster to the metronome’s period at the beginning of each block. Additionally in both halves the plauteau of convergence of the IMI curves is slightly but consistently higher than the metronome’s period. One would expect that in second half the participants should be able to converge more accurately to the period of the metronome. These observations about the IMI alignment reveal that it is very likely that there is no long-term anticipatory mechanism being deployed for the period alignment and that it is most likely an on-the-fly process affected only by the immediately preceding one or two trials. This type of mechanism for period alignment would fit well with the lag-1 or lag-2 models that have been developed to capture the steady-state behavior of sensorimotor synchronization (Michon & van der Valk, 1967; Pressing, 1998; Pressing & Jolley-Rogers, 1997; Schulze & Vorberg, 2002; Semjen et al., 1998; Vorberg, 1996). (Hary & Moore, 1987; Mates, 1994a, 1994b).

Another striking finding of the current study is the fact that the long-term negative gradient of the asynchrony is limb-specific and is not influenced by the interleaved blocks of other limb movements. This is evidence for limb-specific sensorimotor memory. That is, a part of the brain keeps a record of the relative timing between the onset of the previous stimulus (sensory areas) and the corresponding intended movement (motor areas), which record is stored and recalled the next time the same limb is used. Although it is known from previous research that interlimb transfer of acquired motor skills happens, depending in the context of the task(Bao et al., 2022; Yadav & Mutha, 2020), it is the first time to our knowledge that it is demonstrated that basic sensorimotor information such as the asynchrony between visual and motor/tactile parts of the brain is stored and recalled in some form of limb-specific memory.

This observed limb-specificity indicates that some brain areas with somatotopy must be involved and play crucial role in this mechanism. The motor and somatosensory cortices are such areas, which are obviously involved in the brain circuit for sensorimotor synchronization. Another area which has been shown to have somatotopic mapping, containing actually two separate such maps, is the cerebellum(Boillat et al., 2020). The cerebellum together with the basal ganglia form discrete circuits, of reciprocal information flow, with various parts of the cerebral cortex. These segregated circuits are termed “loops”. Each of these circuits serves a role according to the cerebral area it is connected to. Middleton and Strick (Middleton & Strick, 2000) reviewed evidence that there are two such circuits, one loop between cerebellum, basal ganglia, and motor cortex, which is subserving motor actions, and one .loop between cerebellum, basal ganglia and prefrontal cortex, subserving higher cognitive functions. So it would be that the former “motor loop” is responsible in this study for the automatic SMA correction at the beginning of each block, and the short-term, limb-specific somatosensory memory of SMA. And the latter “cognitive loop” is the one performing the IMI alignment, performed in the biggest part of each block.

A last phenomenon that is worth commenting on is that the within-block IMI curve settles consistently towards the end of the block to a level slightly higher than the metronome’s period. Such a bias pertains to the entire experiment although it gets slightly reduced progressively. The source of this bias is unclear. The participants had such a long exposure to this rhythm that in terms of adaptation it would be expected that they would converge to the actual metronome’s period relatively fast. So this bias seems to have a more inherent character. One such bias is known to exist in human behavior, known as Vierordt’s Law (Lejeune & Wearden, 2009). In 1868 Karl Vierordt in his book *Der Zeitsinn nach Versuchen* published the results of a set of experiments, in which he studied with psychophysics the degree of how a perceived time interval is distorted when it is reproduced. The main finding of this time can be summarized as: “*The intervals reproduced are longer than the target time when it is short, but shorter than it when it is long. The* ‘‘*indifference point*’’ *where the reproductions are accurate lies at around 2 (upper panel) or 3 (lower panel) s*.” In the current experiment the metronome period is 1.2 seconds and in this “short” range, according to Vierordt’s Law, it would be expected that participants would reproduce longer IMIs than the metronome’s period.

Vierordt’s Law has been a subject of debate for the last 150 years, with the main criticism being that the observed effect occurs mainly due to the range of durations tested, which according to the “law of central tendency” centers judgement to a center point in the stimulus range below which the stimulus magnitude, in our case duration, is overestimated and above which is underestimated(Hollingworth, 1910). Another strong point of criticism of Vierordt’s Law is that when the randomization used to produce the stimulus sequence has large jumps between successive trials in terms of the employed metronome periods, then this bug differences can create a central tendency such as the one observed in Vierordt’s Law. A recent study (Glasauer & Shi, 2021)tested a “randomized” sequence of durations against a “random-walk” sequence which had eliminated large jumps in the intervals tested. With the “random-walk” sequence the results still had a pattern following Vierordt’s Law but with much smaller levels of over- and underestimation of durations(between +25% to −5%).

In the current experiment we have no range of metronome periods but a single rhythm of 1.2sec, presented in blocks of 10 trials with no stimulus stochasticity between the trials. So there are no effects of the range or sequence of the presented durations. So could the Vierordt’s law still be affecting the estimation of the metronome period in participants’ brains? It could be argued that the brain has already built-in a probability distribution of the temporal intervals encountered in every day life and that this range of values itself is accompanied by a central tendency which creates an effect not as strong as in Vierordt’s experiments but of much smaller magnitude. And this could be the phenomenon manifested here in IMI being consistently larger the the metronome’s period. Other studies have also observed such an overestimation without being able to find a good explanation for it.

In conclusion, the novel findings of this study add significant contributions to understanding sensorimotor synchronization to external rhythms. These are important beyond the limited scope of moving in synchrony to a metronome. They are important for all our actions that are guided by an anticipatory brain mechanism which is tuned to the regularities of the sensory information such as sports and music.

## Notes

### Competing Interest Statement

The authors have declared no competing interest.

